# Collagen I promotes cancer cell survival via amino acid import and mTORC1 activation

**DOI:** 10.1101/2025.11.26.690708

**Authors:** Mona Nazemi, Bian Yanes, Ifeoluwa Oyelade, Heather Walker, Elena Rainero

## Abstract

Invasive breast and pancreatic cancer cells thrive within a collagen I-rich, poorly perfused extracellular matrix (ECM) network, necessitating robust metabolic adaptation to endure nutrient deficiency, such as glucose starvation. Here we demonstrate that collagen I is critical for the survival and growth of breast and pancreatic cancer cells, by maintaining their anabolic status. Mechanistically, collagen I promotes α2β1 integrin-dependent mammalian target of rapamycin complex 1 (mTORC1) activation and drives the membrane localisation of the LAT1−4F2hc amino acid transporter. This process ensures a sustained intracellular essential amino acid supply, further fuelling mTORC1 activity and limiting autophagy. This collagen I-driven pathway is essential for cancer cell survival under nutrient stress, as inhibiting the activity of α2β1 integrin or the LAT1-4F2hc transporter significantly reduces cell growth and invasion in both 2D and 3D models. Finally, the clinical relevance of these transporters is underscored by the significant upregulation of LAT1-4F2hc expression in basal-like breast and pancreatic cancer patients, correlating with poor prognosis and drug resistance. Collectively, our findings highlight that targeting the LAT1−4F2hc transporter might represent a highly promising therapeutic strategy to limit cancer cell growth and invasion in highly fibrotic and nutrient-deprived tumours.

## Introduction

Breast cancer is characterised by a desmoplastic reaction, driven by the accumulation of fibrillar collagens, especially type I ^1^, which is associated with poorer clinical outcomes ^2–4^. In healthy mammary tissue, type I collagen is essential for tissue development and cellular organisation during mammary gland branching ^5–8^. However, upon cancer initiation, type I collagen undergoes structural alterations that significantly impact ECM stiffness and elasticity, with fibres becoming thicker and more aligned ^9–11^. This reorganisation, particularly the radial alignment of collagen fibres, facilitates cancer cell migration and invasion, further promoting metastasis ^10–12^. A similar fibrotic response is observed in pancreatic cancer, where fibrillar collagens, primarily types I and III, comprise over 80% of the ECM in pancreatic ductal adenocarcinoma (PDAC) tumours, where they play a significant role in tumour progression. These collagens provide not only structural support but also act as a reservoir for growth factors and influence the behaviour of both cancer and stromal cells through their organisation, crosslinking, and dynamic changes. Such remodelling can drive key tumorigenic processes including cell proliferation, angiogenesis, invasion, metastasis, resistance to apoptosis, and reactivation from dormancy ^13–15^.

Altered cellular metabolism is a hallmark of cancer. Cancer cell metabolic phenotype is influenced by both cell-intrinsic and extrinsic factors within the tumour microenvironment (TME), including nutrient availability ^16–19^. Unlike normal cells with functional vasculature, cancer cells often grow in poorly perfused environments with limited nutrient access. To survive and sustain high proliferation rates, they reprogram metabolic pathways to optimise nutrient acquisition and utilisation ^20–22^. Central to this reprogramming is the uptake of glucose (Glc) and glutamine, serving as carbon sources and electron donors ^23^. While intrinsic metabolic regulation is increasingly understood, the specific effects of TME-derived nutrient constraints remain poorly characterised.

In this study, we showed that collagen I partially rescued invasive breast cancer and PDAC cell growth under Glc starvation. This was due to a reduction in cell death in breast cancer cells and a combination of increased survival and cell division in PDAC cells. Mechanistically, collagen I promoted α2β1 integrin-dependent mammalian target of rapamycin complex 1 (mTORC1) activation. In addition, collagen I increased the plasma membrane localisation of both the light chain (SLC7A5) and the heavy chain (SLC3A2) of the amino acid transporter LAT1-4F2hc, promoting essential amino acid uptake. Inhibition of either α2β1 integrin or LAT1-4F2hc prevented collagen I-dependent cell growth and invasion under Glc starvation, in both 2D and 3D contexts, including primary mouse tumour organoids. Finally, the co-expression of SLC3A2 and SLC7A5 was significantly upregulated in basal-like breast cancer and pancreatic cancer patients, correlating with poor disease outcomes and drug resistance. Therefore, this work suggests that the inhibition of LAT1-4F2hc could represent a promising therapeutic strategy to limit cancer cell growth and invasion in highly fibrotic and nutrient deprived breast and pancreatic tumours.

## Results

### Collagen I promoted the growth/survival of breast and PDAC cancer cells under glucose starvation

Breast PDAC tumours are characterised by a fibrotic and nutrient deprived TME. To investigate the effect of collagen I on cellular responses to Glc starvation, we cultured breast cancer cells (MDA-MB-231) and pancreatic cancer cells (PANC1) on either 2 mg/ml polymerised collagen I or uncoated plastic under Glc starvation for up to 6 days. We found that the presence of collagen I significantly increased MDA-MB-231 (**Figure 1A**) and PANC1 (**Figure 1B**) cell numbers.

**Figure 1.**
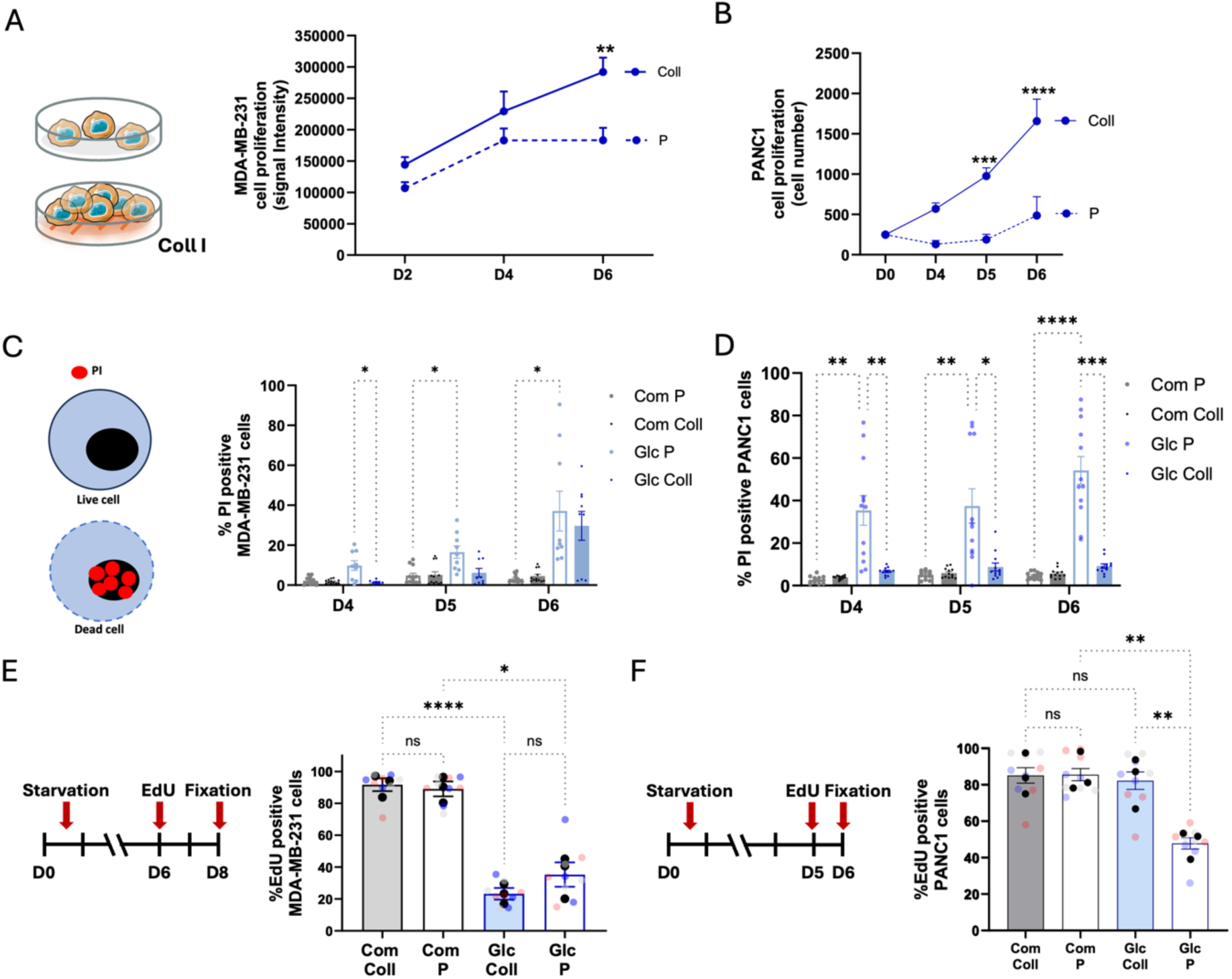
Collagen I supported cell growth under GLc starvation. **(A)** MDA-MB-231 cells were seeded on plastic (P) or 2 mg/ml collagen I (Coll) for 6 days under Glc starvation, fixed, stained with DR, and imaged with a Licor Odyssey Sa system. Signal intensity was calculated by Innage Studio Lite software. (B) PANC1 cells were seeded on 2 mg/ml collagen I (Coll) or plastic (P) under Glc starvation for 6 days, fixed and stained with Hoechst 33342. Images were collected by ImageXpress micro and analysed by MetaXpress software. Data are presented as mean ± SEM, N=3 independent experiments, **p < 0.01, ***p < 0.001, ***p < 0.0001 two-way ANOVA, Tukey’s multiple comparisons test. **(C-D)** MDA-MB-231 and PANC1 cells were seeded or plastic (P) or 2 mg/ml collagen I (Coll) for 6 days under glucose starvation (Glc) or in complete media (Com), every two days cells were treated with PI and Hoechst 33342 for 1hr, imaged with ImageXpress micro and analysed by MetaXpress software. Data are presented as mean ± SEM, N=3 independent experiments, *p<0.05, **p < 0.01, ***p < 0.001, ****p < 0.0001 two-way ANOVA, Tukey’s multiple comparisons test. (E-F) MDA-MB-231 and PANC1 cells were seeded on 2 mg/ml collagen I (Coll) or plastic (P), in complete media (Com) or under Glc starvation (Glc). MDA-MB-231 cells were incubated with EdU at day 6 post starvation, fixed and stained with Hoechst 33342 and Click iT EdU imaging kit at day 8. PANC1 cells were incubated with EdU at day 5 post starvation, fixed and stained with Hoechst 33342 and Click iT EdU imaging kit at day 6. Images were collected by ImageXpress micro and analysed by MetaXpress software. Data are presented as mean ± SEM, N=3 independent experiments (black dots represent the mean of individual experiments), *p < 0.05, **p < 0.01, ****p < 0.0001 one-way ANOVA Kruskal-Wallis, Dunn’s multiple comparisons test.

To determine whether collagen I-driven growth was associated with carcinoma progression, we took advantage of the MCF10 series of cell lines, composed of non-transformed mammary epithelial (MCF10A), non-invasive ductal carcinoma in situ (MCF10A-DCIS) and invasive (MCF10CA1) cells. In contrast to MDA-MB-231 cells, collagen I did not support the growth of MCF10A (**Figure S1A**) and MCF10A-DCIS (**Figure S1B**) cells under Glc starvation. However, collagen I promoted the growth of MCF10CA1 cells after 6 days of Glc deprivation (**Figure S1C**). Consistently, we obtained similar results in SW1990 pancreatic cancer cells, where the presence of collagen I significantly increased cell numbers compared to plastic (**Figure S2A**). Together, these data indicate that collagen I-mediated cell growth under Glc starvation is associated with carcinoma progression.

To elucidate whether the observed increase in cell numbers was due to enhanced survival or increased division rates, we performed cell death and EdU incorporation assays. Propidium Iodide (PI) is a cell-impermeable nuclear dye, which is excluded from healthy cells, while can penetrate damaged or dying cells. The quantification of the percentage of PI-positive cells indicated that MDA-MB-231 cells exhibited a higher death rate on plastic under Glc starvation compared to complete media at all time points. Moreover, collagen I significantly reduced cell death under Glc starvation at day 4, with a similar trend at day 5, although not statistically significant (**Figure 1C**). Similarly, PANC1 cell death was significantly higher under Glc starvation compared to complete media on plastic, while the presence of collagen I completely prevented cell death at all time points (**Figure 1D**). Collagen I did not affect viability in both MDA-MB-231 and PANC1 cells in complete media (**Figure 1C,D**). 5-ethynyl-2′-deoxyuridine (EdU) is a thymidine analogue that is incorporated into the DNA during the S phase of the cell cycle. As expected, we detected a significant reduction in the percentage of EdU positive cells under Glc starvation compared to complete media on plastic in both MDA-MB-231 and PANC1 cells (**Figure 1E,F**). While there was no significant difference in EdU incorporation in MDA-MB-231 cells in the presence or absence of collagen I under Glc deficiency (**Figure 1E**), there was a significantly higher percentage of EdU-positive PANC1 cells cultured on collagen I (**Figure 1F**), indicating an increased division rate. Collagen I did not affect EdU incorporation in complete media in either cell line (**Figure 1E,F**). Moreover, the presence of collagen I significantly increased MCF10CA1 EdU incorporation (**Figure S1D**) and reduced the percentage of MCF10CA1 cells positive for the apoptosis marker cleaved caspase 3/7 (**Figure S1E**). Similarly, collagen I promoted EdU incorporation (**Figure S2B**) and reduced cell death in SW1990 cells (**Figure S2C**).

In summary, our findings demonstrate that the presence of collagen I increased MDA-MB-231, MCF10CA1, SW1990 and PANC1 cell numbers under Glc starvation. This effect was attributed to a reduction in cell death for all cell lines, with a concomitant increase in the division rate in PANC1, MCF10CA1 and SW1990 cells, but not in MDA-MB-231 cells. Moreover, the observation that collagen I did not affect the growth of non-invasive cancer and normal mammary epithelial cells suggests that this survival mechanism is associated with invasiveness and/or cancer progression.

### α2β1 integrin was required for collagen I-dependent cell growth

We have previously demonstrated that ECM internalisation supported breast cancer cell growth under amino acid starvation ^24^. To investigate whether collagen I-dependent cell growth/survival under Glc starvation was also mediated by its internalisation and subsequent lysosomal degradation, we assessed the ability of MDA-MB-231 cells to uptake collagen I under starvation. Interestingly, we found that collagen I endocytosis was significantly increased in Glc-depleted media, both in the presence and absence of the lysosomal protease inhibitor E64d (**Figure S3**). We have previously showed that glutaraldehyde-mediated cross-linking prevents ECM extracellular degradation and internalisation ^24^. To determine the requirement of collagen I uptake in promoting cell growth, MDA-MB-231 and PANC1 cells were seeded on chemically cross-linked collagen I. While MDA-MB-231 cells exhibited comparable growth on cross-linked collagen I as on control collagen I (**Figure 2A**), cross-linking further stimulated PANC1 cell growth (**Figure 2B**), indicating that the supportive effect of collagen I was not primarily dependent on its internalisation.

**Figure 2.**
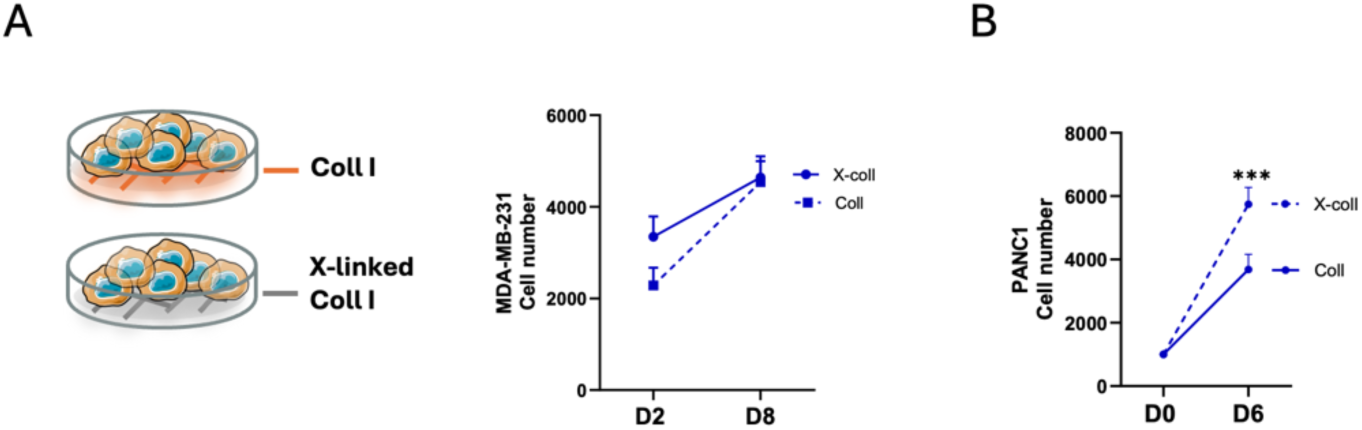
Collagen I chemical crosslinking did not impair cell proliferation under starvation. MDA-MB-231 **(A)** and PANC1 cells **(B)** were seeded on 2 mg/ml collagen I (Coll) or cross-linked collagen I (X-coll) for 8 or 6 days, respectively. Cells were fixed and stained with Hoechst 33342. Images were collected by an ImageXpress micro and analysed by MetaXpress software. Data are presented as mean ± SEM, N=3 independent experiments, ***p < 0.001 two-way ANOVA, Tukey’s multiple comparisons test.

Cells interact with the ECM through plasma membrane receptors of the integrin family, including α2β1 integrin, the main collagen I receptor ^25^. Upon ligand binding, integrins trigger the activation of adhesion signalling pathways, which promote cell proliferation and survival ^26^. Therefore, we examined the role of α2β1 by treating the cells with BTT-3033, an inhibitor that prevents α2β1 collagen binding, and monitoring cell proliferation. BTT-3033 treatment elicited a dose-dependent inhibition on MDA-MB-231 and PANC1 cell proliferation (**Figure 3A**). Consistently, α2 integrin downregulation significantly decreased cell numbers in both cell lines. In contrast, β1 knockdown did not affect MDA-MB-231 cells but significantly reduced PANC1 cell numbers (**Figure 3B**). Immunofluorescence staining confirmed efficient downregulation of both α2 and β1 integrin protein levels in both cell lines (**Figure S4**). Similarly, both pharmacological inhibition and siRNA-mediated knock down of α2 integrin significantly impaired MCF10CA1 cell growth on collagen I under Glc starvation (**Figure S5**). Furthermore, to extend our findings to a more physiologically relevant context, we evaluated the impact of BTT-3033 and α2 integrin knockdown in 3D models. 3D spheroids were generated, embedded in a mixture of collagen I and Geltrex and grown under Glc starvation for 2 days (MDA-MB-231 cells) or 6 days (PANC1 cells). MDA-MB-231 spheroids were highly invasive and extended long multicellular protrusions. Both α2 pharmacological inhibition and siRNA-mediated protein downregulation significantly inhibited MDA-MB-231 spheroid invasion (**Figure 3C,D**). In contrast, PANC1 spheroids mostly grew in size, with some cells invading in the surrounding ECM. Consistently, α2 inhibition and knockdown dampened spheroid growth under Glc starvation (**Figure 3E,F**). Moreover, primary tumour organoids were generated by growing E0771 mouse triple negative breast tumours into a mixture of collagen I and Geltrex in Glc-free media, in the presence or absence of BTT-3033 for 3 days. While control organoids grew over time, with strands of cells invading the surrounding ECM, both 2.5 μM and 5 μM BTT-3033 completely prevented organoid growth and opposed cell invasion. Image analysis indicated that α2 pharmacological inhibition significantly reduced organoids’ size and invasion compared to the control group (**Figure 3G,H**). These results collectively indicate that the collagen I receptor α2β1 integrin modulated collagen I-dependent cell growth under Glc starvation in breast and pancreatic cancer cells, in both 2D and 3D contexts.

**Figure 3.**
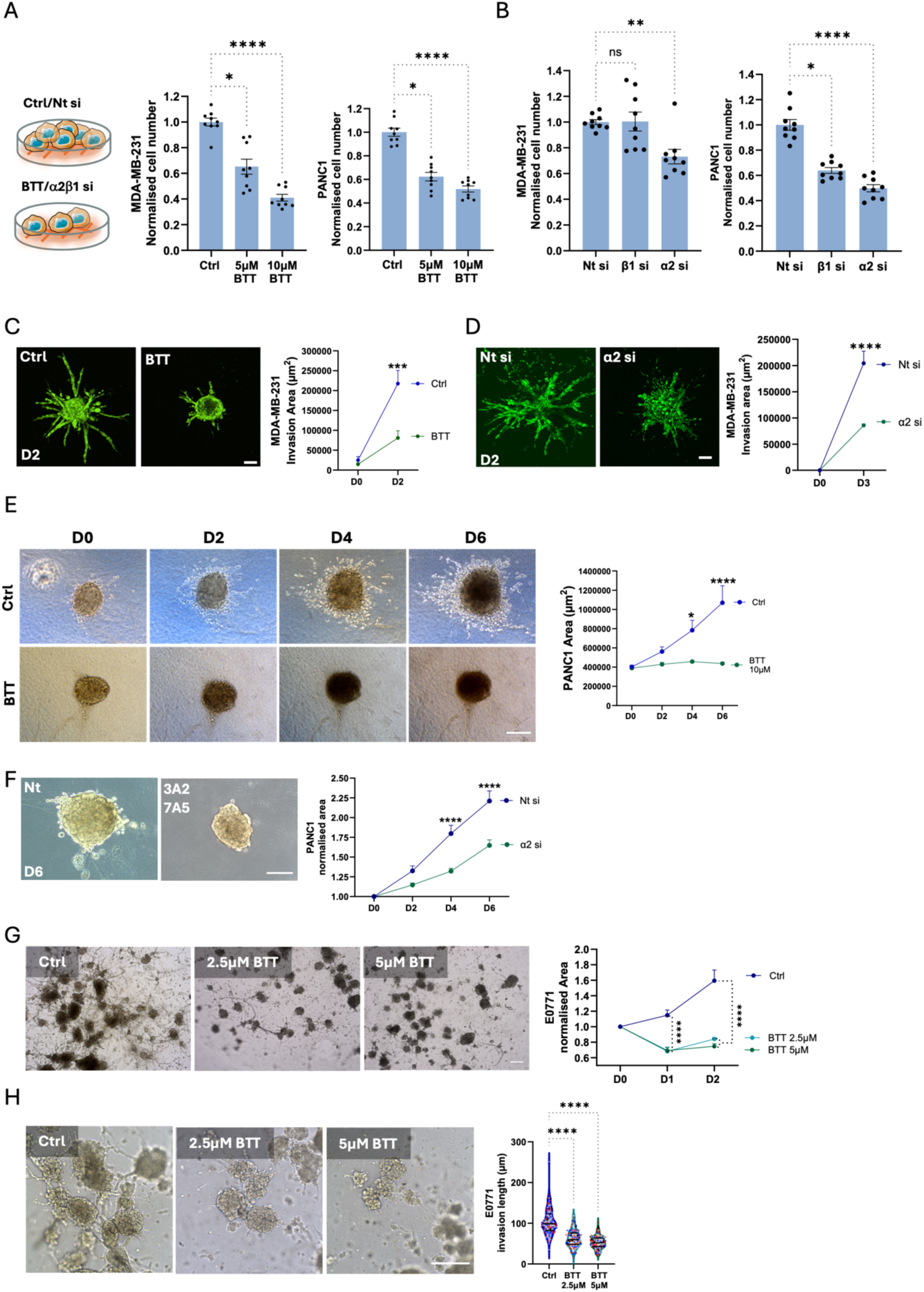
α2β1 integrin was required for cancer cell growth and invasion under Glc starvation. **(A)** MDA-MB-231 and PANC1 cells were seeded on 2 mg/ml collagen I and treated with 5 µM, 10 µM BTT-3033 (BTT) or DMSO (Ctrl) for 4 days under Glc starvation. (B) MDA-MB-231 and PANC1 cells were seeded on 2 mg/ml collagen I and transfected with an siRNA targeting β1 integrin (β1 si), an siRNA targeting α2 integrin (α2 si) or a non-targeting siRNA control (Nt si) for 4 days under Glc starvation. Cells were fixed and stained with Hoechst 33342. Images were collected by ImageXpress micro and analysed by MetaXpress software. (C) MDA-MB-231-GFP spheroids were generated by the hanging drop method, embedded in 3 mg/ml collagen I and Geltrex (50:50) mixture and starved in Glc-free media for 2 days, in the presence of 10 µM BTT-3033 (BTT) or DMSO (Ctrl). Live images were collected by a Nikon A1 Confocal microscope. **(D)** MDA-MB-231-GFP cells were transfected with an siRNA targeting α2 integrin (α2 si) or a non-targeting siRNA control (Nt si) for 24 hours. Spheroids were generated by the hanging drop method, embedded in 3 mg/ml collagen I and Geltrex (50:50) mixture in Glc starvation media for 3 days. Live images were collected by a Nikon A1 Confocal microscope. (E) PANC1 spheroids were generated by the hanging drop method, embedded in 3 mg/ml collagen I and Geltrex (50:50) mixture in Glc starvation media for 6 days in the presence of 10 µM BTT-3033 (BTT) or DMSO (Ctrl). Spheroids were imaged live every 2 days with an Olympus E45O microscope. (F) PANC1 cells were transfected with an siRNA targeting α2 integrin (α2 si) or a non-targeting siRNA control (Nt si) for 24 hours. Spheroids were generated by the hanging drop method, embedded in 3 mg/ml collagen I and Geltrex (50:50) mixture in Glc starvation media for 6 days. Live images were collected by an Olympus E45O microscope. **(G, H)** E0771 mouse tumour organoids were grown in a 3 mg/ml collagen I and Geltrex (50:50) mixture in Glc starvation media for 3 days in the presence of 2.5 µM, 5 µM BTT-3033 (BTT) or DMSO (Ctrl). Spheroids were imaged live every day with an Olympus E45O microscope. Bar, 250 µm. Data are presented as mean ± SEM, N=3 independent experiments, *p < 0.05, **p < 0.01, ***p < 0.001, ****p < 0.0001; (A, B and H) Kruskal-Wallis, Dunn’s multiple comparisons test; (C-G) Two-way ANOVA, Tukey’s multiple comparisons test.

### Collagen I modulated mTOR signalling and autophagy

It is well established that Glc controls nutrient signalling, through the regulation of mTORC1 and AMP-activated protein kinase (AMPK). Under Glc starvation, mTORC1 activity is inhibited, leading to the activation of autophagy, an intracellular degradation process to support nutrient acquisition. This is a crucial adaptation where cells shift from growth and anabolic processes promoted by active mTORC1 signalling to autophagy-mediated catabolism for survival and nutrient recycling ^27^. To determine if the increased cell numbers and survival rates observed on collagen I under Glc deficiency were mediated by changes in nutrient signalling, we assessed mTORC1 signalling and autophagy. As a readout for mTORC1 activation, we stained the cells for the phosphorylated form of S6 (pS6), a well-established mTORC1 downstream target. Surprisingly, we did not detect changes in S6 phosphorylation in response to one-day Glc starvation compared to complete media, while collagen I consistently increased the phosphorylation of S6 in MDA-MB-231 cells, regardless of nutrient availability (**Figure 4A**). BTT-3033 treatment significantly lowered S6 phosphorylation in the presence of collagen I to the same level as in cells seeded on plastic (**Figure 4A**), indicating that α2 integrin was required for collagen I-driven activation of mTORC1 signalling. Interestingly, the phosphorylation of another mTORC1 downstream target, 4EBP1, was significantly reduced under Glc starvation, independently on the substrate the cells were seeded on (**Figure S6**), suggesting that adhesion signalling might affect only a subset of mTORC1 targets. Consistently, S6, but not 4EBP1, phosphorylation has been reported to be associated with focal adhesions ^28^.

**Figure 4.**
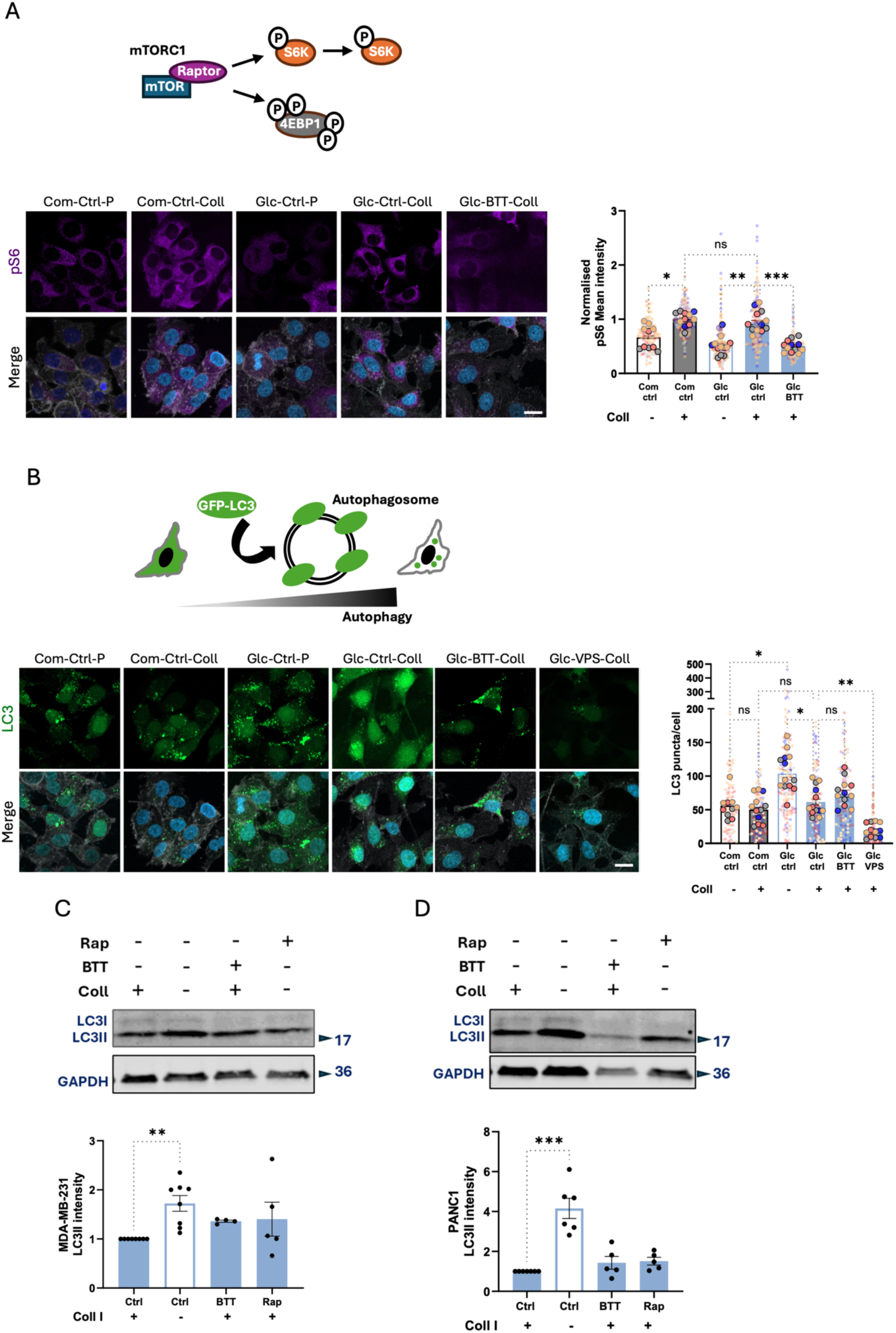
Collagen I promoted mT0RC1 activation and inhibited autophagy in cancer cells. **(A)** MDA-MB-231 cells were seeded on 2 mg/ml collagen I (Coll, +) or plastic (P, -) under complete or Glc starvation media for one day, in the presence of 10 µM BTT-3033 (BTT) or DMSO (Ctrl). Cells were fixed and stained with Hoechst 33342 (blue), Phalloidin (grey) and pS6 (magenta). (B) MDA-MB-231 cells stably expressing GFP-LC3 were seeded as in A, in the presence of 4 µM VPS34-IN1 (VPS) where indicated. Cells were fixed and stained with Hoechst 33342 (blue) and Phalloidin (grey). Images were collected with a ZEISS LSM98O Airyscan2 confocal microscope and analysed by Fiji/lmageJ software. Bar, 20 µm. Data are presented as mean ± SEM, N=3 independent experiments (the bigger dots represent the mean of individual fields of view). MDA-MB-231 (C) and PANC1 (D) cells were seeded on 2 mg/ml collagen I (+) or plastic (-) under Glc starvation. Cells were treated with DMSO (Ctrl), 100 nM rapamycin (Rap) or 10 µM BTT-3033 (BTT). Cell lysates were collected after one (C) or three days (D) of starvation. Data are presented as mean ± SEM, N>4 independent experiments, *p < 0.05, **p < 0.01, ***p < 0.001, Kruskal-Wallis, Dunn’s multiple comparisons test.

To investigate whether collagen I controlled autophagy, we used MDA-MB-231 cells stably expressing a GFP-LC3 reporter. During autophagy, microtubule-associated protein 1A/1B-light chain 3 (LC3) is converted from a soluble form (LC3-I) to a membrane bound form (LC3-II) which localises to the membrane of autophagosomes, double-membrane vesicles that engulf cellular components ^29^. Therefore, if autophagy is induced, GFP-LC3 is recruited on autophagosomes, resulting in a punctate distribution, while when autophagy is oG, GFP-LC3 is mostly diffused in the cytosol. As anticipated, one-day Glc starvation on plastic significantly increased the number of LC3-positive puncta, indicating an induction of autophagy. Consistent with the fact that ECM attachment prevents autophagy ^30^, collagen I reduced the number of LC3-positive structures under Glc starvation, to the same levels as cells in complete media (**Figure 4B**). The presence of BTT-3033 resulted in a small, but not statistically significant, increase in LC3 puncta, indicating that α2 integrin-mediated adhesion did not mediate collagen I-dependent autophagy inhibition under Glc starvation. As expected, the inhibition of VPS34, a known autophagy regulator, significantly reduced GFP-LC3 puncta (**Figure 4B**). The decrease in LC3 puncta in the presence of collagen I could represent an increase in autophagic flux, where autophagosomes are being degraded faster, or a decrease in autophagosome formation. To address this, we used the LC3-RFP-GFP reporter to measure autophagic flux, based on the different sensitivities of its fluorescent proteins to lysosomal acidity. GFP fluorescence is quenched at acidic lysosomal pH, while RFP remains stable. When autophagosomes fuse with lysosomes, the GFP signal decreases faster than RFP, making the RFP/GFP ratio an indicator of autophagic activity ^31^. The stable ratio of LC3-RFP (autolysosome) to LC3-Yellow (autophagosome) between plastic and collagen I suggested that the increased LC3 vesicular recruitment in plastic-cultured cells was not due to disrupted lysosomal degradation, nor the reduction in LC3 puncta on collagen I was caused by increased degradation (**Figure S7**). Consistently, western blot analysis of LC3-II protein levels in MDA-MB-231 and PANC1 cells showed that cells cultured on plastic exhibited significantly higher LC3-II levels compared to those on collagen I after one or three-day Glc starvation, respectively. Similarly to our imaging results, inhibition of α2β1 integrin did not significantly affect LC3-II levels. Moreover, the mTORC1 inhibitor Rapamycin also did not influence LC3-II, suggesting that the reduction in autophagy observed in the presence of collagen I was not mediated by increased mTORC1 activity (**Figure 4C,D**).

Collectively, our data indicate that cells cultured on collagen I under Glc starvation maintain an anabolic status, characterised by higher mTORC1 activation and lower autophagy, whereas cells on plastic are driven towards catabolism. Interestingly, α2β1 integrin is required for mTORC1 activation, but not for autophagy inhibition, suggesting alternative regulatory mechanisms. This implies that the presence of collagen I alleviates the metabolic stress induced by Glc deprivation, enabling the cells to maintain an anabolic state.

### Collagen I increased intracellular amino acid content

It is well established that cancer cells rewire their metabolism under Glc starvation, by increasing their reliance on alternative fuels, such as amino acids ^27^. In addition, several amino acids promote mTORC1 activity ^32^. Therefore, we hypothesised that collagen I might result in changes in intracellular metabolite levels. To address this, we performed non-targeted mass spectrometry in cells grown under Glc starvation for one day (**Figure 5A,D**) and detected significant metabolic differences in MDA-MB-231, PANC1, MCF10CA1 and SW1990 cells cultured on collagen I compared to plastic (**Figure 5B,E** and **Figure S8A,B,D**). Furthermore, MDA-MB-231 cells grown on chemically cross-linked collagen I showed no significant difference in metabolic content relative to untreated collagen I (**Figure S9**), indicating that collagen I internalisation was not responsible for the collagen I-induced changes in intracellular metabolites. Metabolic pathway enrichment analysis on the upregulated metabolites on collagen I identified several pathways related to amino acid metabolism in MDA-MB-231 and PANC1 cells (**Figure 5C,F**), as well as in MCF10CA1 and SW1990 cells (**Figure S8C,E**), suggesting increased intracellular amino acid levels. To confirm this, we conducted targeted mass spectrometry during a four-day Glc starvation period. Overall, the levels of most amino acids were increased on collagen I compared to plastic. In particular, essential amino acids, including leucine/isoleucine, lysine, methionine, phenylalanine, threonine, tryptophan, and valine, were upregulated on collagen I in MDA-MB-231 and PANC1 cells, from day one to day four of starvation (**Figure 5G,H**). To determine the source of the upregulated metabolites, we inhibited autophagy by treating MDA-MB-231 and PANC1 cells with a VPS34 inhibitor. Most amino acid levels were unaffected by the VPS34 inhibitor, except for the following non-essential amino acids: aspartate, glycine, and glutamine in MDA-MB-231 cells (**Figure 5G**), and alanine, aspartate, glycine, glutamine, and serine in PANC1 cells (**Figure 5H**). Interestingly, these amino acids were also the most abundant in cells on plastic, consistent with our observation that autophagy was elevated on plastic compared to collagen I. Therefore, we ruled out autophagy as the main source for elevated amino acid levels on collagen I. Given that all upregulated amino acids are essential and present in the culture media, we hypothesised that the cells might rely on extracellular amino acid import.

**Figure 5.**
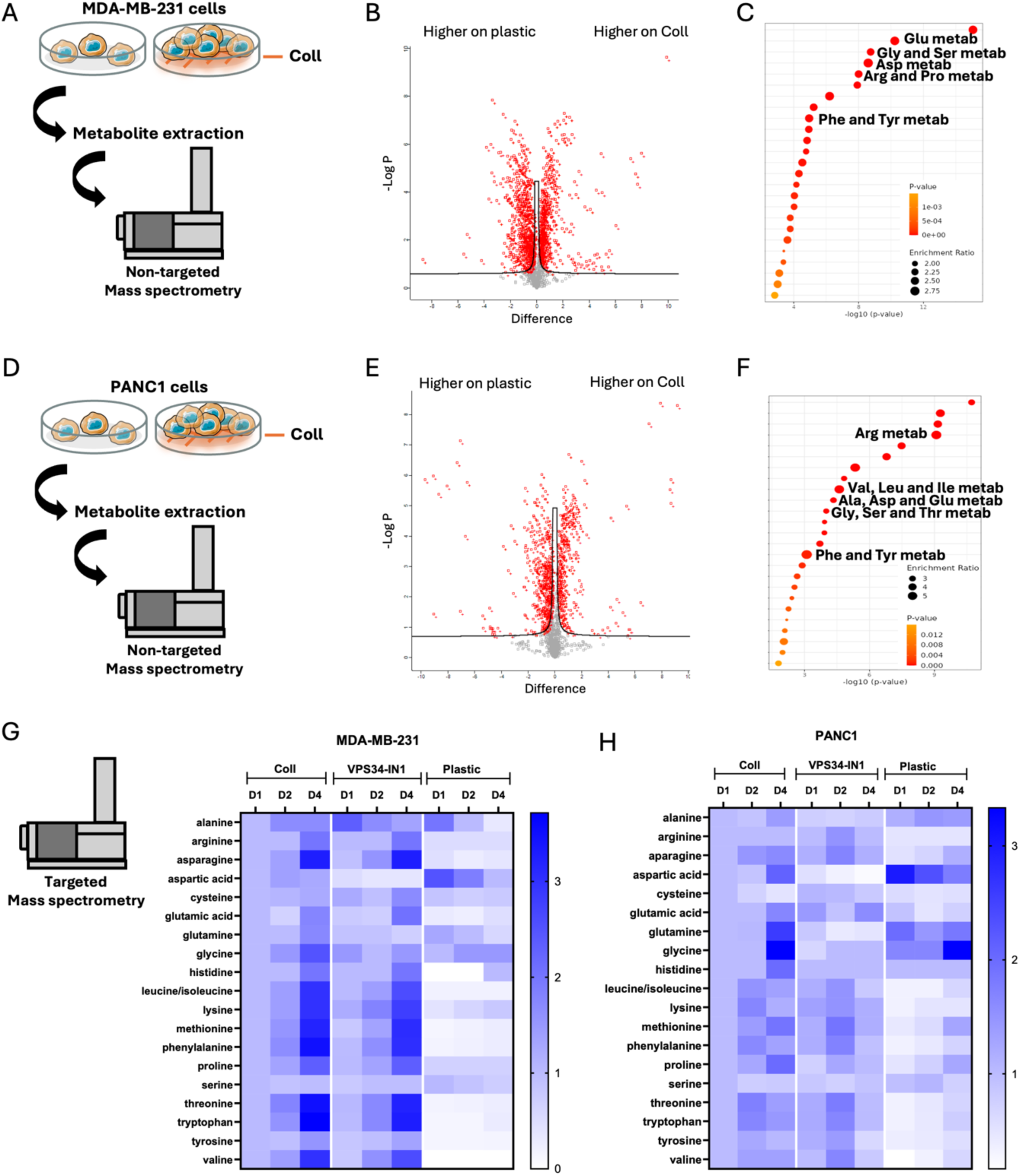
Collagen I increased intracellular amino acid levels. **(A,D)** Metabolomics workflow, MDA-MB-231 and PANC1 cells were plated on plastic or 2 mg/ml collagen I (coll) for one day in Glc starvation media. Metabolites were extracted and quantified by non-targeted mass spectrometry. Volcano plot (B,E) and enriched metabolic pathways **(C,F)** are shown. **(G)** MDA-MB-231 and **(H)** PANC1 cells were plated on plastic or 2 mg/ml collagen I for up to four days in Glc starvation media in the presence of 4 µM VPS34-IN1 where indicated. Metabolites were extracted at day 1,2 and 4 and intracellular amino acid levels were quantified by targeted mass spectrometry.

### Collagen I increased LAT1-4F2hc protein levels to promote essential amino acid uptake

The majority of the upregulated amino acids (leucine/isoleucine, methionine, phenylalanine, threonine, tryptophan, and valine), excluding threonine, are known substrates for the Large Neutral Amino Acid Transporter 1 (LAT1). LAT1-4F2hc is a heterodimer consisting of the common heavy chain, SLC3A2, and its specific light chain, SLC7A5 (**Figure 6A**). To explore whether LAT1-4F2hc contributed to collagen I-dependent amino acid import, we measured the expression of SLC3A2 and its associated light chains, SLC7A5, SLC7A6, and SLC7A8. qPCR analysis showed that the mRNA levels of SLC3A2 and SLC7A5 were strongly elevated by Glc starvation on plastic and collagen I in both MDA-MB-231 and PANC1 cells, while SLC7A6 and SLC7A8 expression was overall lower and not affected by the starvation (**Figure 6B,C**). We then assessed SLC3A2 and SLC7A5 membrane distribution by performing immunofluorescence staining of non-permeabilised cells. We observed significantly higher SLC3A2 and SLC7A5 protein levels in both MDA-MB-231 (**Figure 6D,E**) and PANC1 (**Figure 6F,G**) cells grown on collagen I compared to plastic. These results collectively suggest that collagen I controls LAT1-4F2hc membrane localisation, rather than its mRNA expression, thereby promoting amino acid import. Indeed, siRNA-mediated knockdown of SLC3A2 and SLC7A5 alone or in combination resulted in lower threonine, phenylalanine, valine, leucine/isoleucine, tryptophan and tyrosine level in MDA-MB-231, as detected by targeted mass spectrometry (**Figure 6H**). In PANC1 cells, the combined downregulation of SLC3A2 and SLC7A5 reduced the intracellular levels of all the amino acid tested (**Figure 6I**). Together these data indicate that collagen I increased intracellular amino acid levels by promoting LAT1-4F2hc-mediated import.

**Figure 6.**
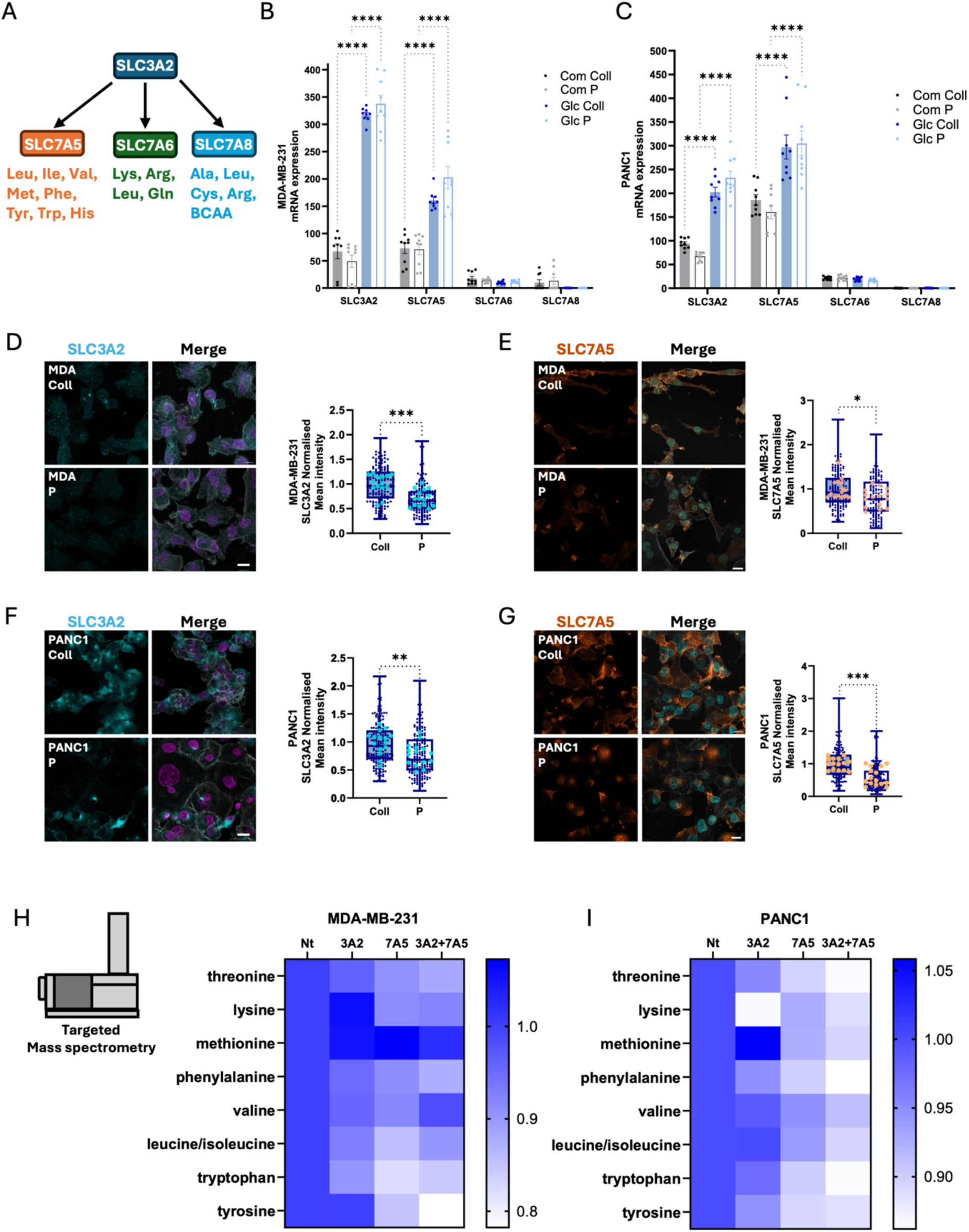
Collagen I promoted LAT1-4F2hc-mediated amino acid import. **(A)** Schematic representation of SLC3A2-containing amino acid transporters. **(B)** MDA-MB-231 and **(C)** PANC1 cells were seeded on 2 mg/ml collagen I (Coll) or plastic (P) under complete or Glc starvation media for 24 hours, mRNA was extracted and the levels of SLC3A2, SLC7A5, SLC7A6, SLC7A8 and GAPDH as housekeeping gene were quantified by SYBR-green qPCR. Data are presented as 2^-ΔCT^; mean ± SEM, N=3 independent experiments, ****p < 0.0001 Kruskal-Wallis, Dunn’s multiple comparisons test. **(D,E)** MDA-MB-231 and **(F,G)** PANC1 cells were seeded on 2 mg/ml collagen I (Coll) or plastic (P) under Glc starvation for one or two days, respectively. Cells were fixed and stained with (D,F) Hoechst 33342 (magenta), SLC3A2 (cyan) and Phalloidin Alexa Fluor 555 (grey) or (E,G) Hoechst 33342 (cyan), SLC7A5 (red) and Phalloidin Alexa Fluor 555 (grey). Bar, 20 µm. Images were collected with a ZEISS LSM98O Airyscan2 microscope and analysed by Fiji/lmageJ software. Data are presented as mean ± SEM, N=3 independent experiments (the bigger dots represent mean intensity of each field of view); **p < 0.01, ***p < 0.001 Mann-Whitney test. (H, I) MDA-MB-231 and PANC1 cells were transfected with an siRNA targeting SLC3A2 (3A2), an siRNA targeting SLC7A5 (7A5) or a combination of siRNA targeting SLC3A2 and SLC7A5 (3A2+7A5) and grown on 2mg/ml collagen I under Glc starvation for one or three days, respectively. Metabolites were extracted and analysed by targeted mass spectrometry. N=3 independent experiments.

### LAT1-4F2hc supported mTORC1 activation and reduced autophagy under glucose starvation, to promote cell growth

As intracellular amino acids are well defined regulators of mTORC1 activity ^32^, we monitored S6 phosphorylation and autophagy upon SLC3A2 downregulation (**Figure 7A,F,K**). We found that SLC3A2 knockdown significantly reduced mTORC1 activity, to a similar extent than upon α2 integrin downregulation (**Figure 7B,G,L**), and enhanced LC3-II levels (**Figure 7C,H,M**) in MDA-MB-231 cells after 1 day of Glc starvation, as well as in PANC1 cells after 2 and 3 days of Glc starvation. Consistent with our pharmacological inhibitor results, α2 integrin knockdown did not affect LC3-II levels (**Figure 7C,H,M**). Moreover, α2 integrin and SLC3A2 downregulation did not influence each other levels, while the siRNA-mediated knockdown significantly reduced the expression of the target proteins (**Figure 7D,E,I,J,N,O**). This suggests that the LAT1-4F2hc transporter mediated collagen I-dependent mTORC1 activation and autophagy inhibition in cancer cells under Glc starvation.

**Figure 7.**
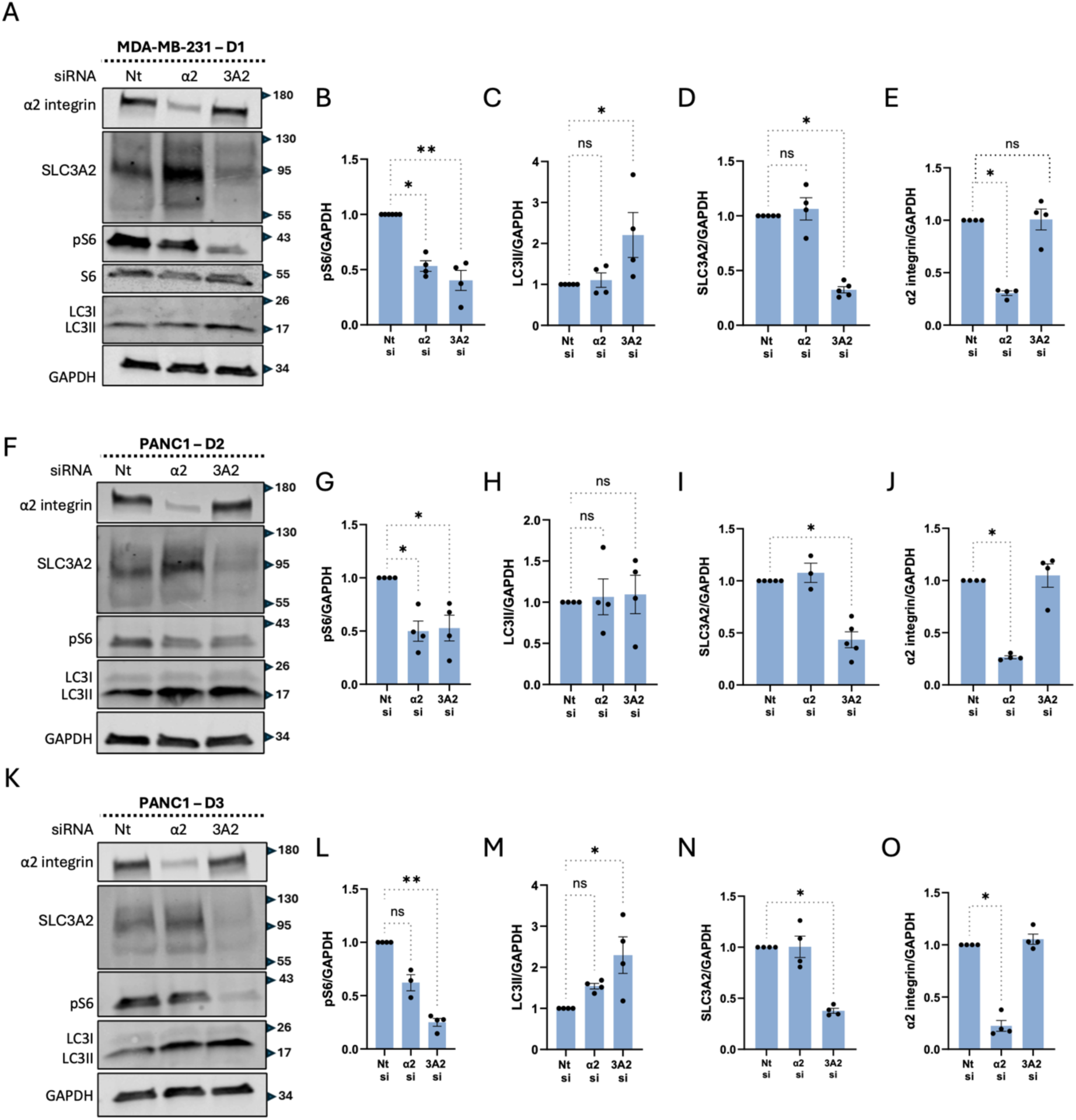
SLC3A2 was required for collagen l-induced mTORCI activation and autophagy inhibition. MDA-MB-231 **(A-E)** and PANC1 **(F-O)** cells were transfected with an siRNA targeting SLC3A2 (3A2 si), an siRNA targeting α2 integrin (α2 si) or a non-targeting siRNA control (Nt si) and grown on 2 mg/ml collagen I in Glc starvation media for one (A-E), two (F-J) and three days (K-O). Protein level of phosphorylated S6 (pS6, B,G,L), LC3II (C,H,M), SLC3A2 (D,I,N) and α2 integrin (E,J,O) were measured via Western blotting. Data are presented as mean ± SEM, N≥4 independent experiments, *p < 0.05, **p < 0.01 Kruskal-Wallis, Dunn’s multiple comparisons test.

We therefore wanted to investigate the role of SLC3A2 and SLC7A5 in supporting cell growth/survival on collagen I. In MDA-MB-231 cells, on the one hand, siRNA-mediated knockdown of SLC3A2 significantly decreased cell numbers under Glc starvation (**Figure 8A**), but not in complete media (**Figure 8B**). On the other hand, SLC7A5 downregulation resulted in a small, but statistically significant, reduction in cell numbers under both Glc starvation (**Figure 8C**) and complete media (**Figure 8D**). In PANC1 cells, SLC3A2 knockdown significantly reduced cell numbers in complete media (**Figure 8F**), but not under Glc starvation (**Figure 8E**). In contrast, SLC7A5 downregulation significantly reduced cell numbers under both conditions (**Figure 8G,H**), indicating an SLC7A5 broader role in PANC1 cell proliferation, regardless of nutrient availability. In the 3D spheroid model, SLC3A2 and SLC7A5 double knockdown resulted in a small, but statistically significant, reduction in MDA-MB-231 spheroid invasion after 3 days of Glc starvation (**Figure 8I**). Additionally, SLC3A2 and SLC7A5 double knockdown reduced PANC1 spheroid size after 6 days of Glc starvation (**Figure 8J**). To determine the role of LAT1-4F2hc in primary tumour cells, E0771 mouse triple negative breast organoids were grown into a mixture of collagen I and Geltrex in Glc-free media in the presence or absence of D-phenylalanine for 2 days. D-phenylalanine is established to block LAT1-4F2hc mediated amino acid transport ^33, 34^. Consistent with our 3D spheroid results, LAT1-4F2hc pharmacological inhibition significantly reduced organoids’ growth and invasion compared to the control group (**Figure 8K,L**). Together, these results demonstrate that SLC3A2 and SLC7A5 are required for breast and pancreatic cell proliferation and invasion, in both 2D and 3D models, to a different extent depending on the cell type.

**Figure 8.**
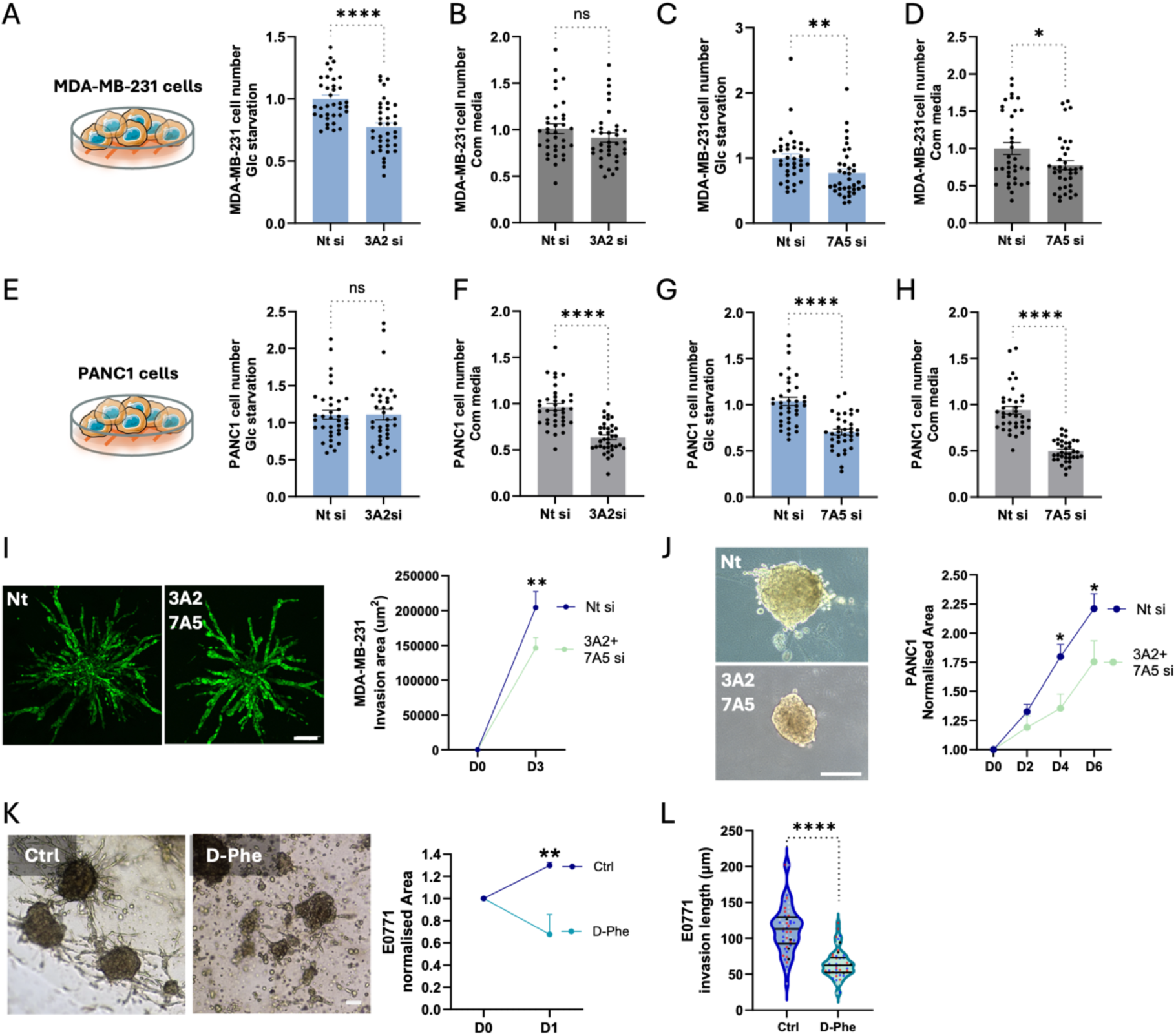
SLC3A2 and SLC7A5 were required for collagen l-dependent celL growth in 2D and 3D systems. MDA-MB-231 **(A-D)** arid PANC1 (E-H) were transfected with an siRNA targeting SLC3A2 (3A2 si), an siRNA targeting SLC7A5 (7A5 si) or a non-targeting siRNA control (Nt si) and grown on 2 mg/ml collagen I under complete or Glc starvation media for 4 days. Cells were fixed and stained with Hoechst 33342. Images were collected by an ImageXpress micro and analysed by Meta×press software. Data are presented as mean ± SEM, N=3 independent experiments, *p < 0.05, **p < 0.01 and ****p < 0.0001 Mann-Whitney test. (I) MDA-MB-231-GFP cells were transfected with a combination of siRNAs targeting SLC3A2 and SLC7A5 (3A2+7A5 si) or a non­targeting siRNA control (Nt si) for 24 hours, spheroids were generated by the hanging drop method and embedded in 3 mg/ml collagen I and Geltrex (50:50 ratio) for 3 days under Glc starvation. Images were collected by a Nikon Confocal A1 microscope. Invasion area was quantified with Fiji/lmageJ. Bar, 200 µm. Data are presented as mean ± SEM, N=2 independent experiments. (J) PANC1 cells were transfected as in I, spheroids were generated by the hanging drop method and embedded in 3 mg/ml collagen land Geltrex (50:50 ratio) for 6 days under Glc starvation. Live images were collected every day by an Olympus E45O microscope. Bar, 250 µm. Data are presented as mean ± SEM, N=2 independent experiments. **(K, L)** E0771 mouse tumours organoids were grown in a 3 mg/ml collagen I and Geltrex (50:50) mixture and starved in Glc starvation media for 2 days in the presence or absence of 50 mM D-phenylalanine (D-Phe). Spheroids were imaged live every day by an Olympus E45O microscope. Bar, 250 µm. Data are presented as mean ± SEM, N=3 independent experiments, *p < 0.05, **p < 0.01 and ****p < 0.0001; (l,J,K) Two-way ANOVA, Tukey’s multiple comparisons test; (L) Mann-Whitney test.

### LAT1-4F2hc elevated expression correlated with poor prognosis and reduced chemotherapy response in breast and pancreatic cancer patients

To examine whether LAT1-4F2hc may have clinical implications, we looked at the relationship between the co-expression of SLC3A2 and SLC7A5 and disease outcome in breast and pancreatic cancer patients. RNA sequencing data showed overexpression of SLC3A2 and SLC7A5 in basal-like breast cancer (**Figure 9A**) and pancreatic adenocarcinoma patients (**Figure 9C**). Correspondingly, concomitant high expression of both genes correlated with poor overall survival (**Figure 9B**) and disease-free survival (**Figure 9D**) in basal-like breast and pancreatic cancer patients, respectively. To determine whether essential amino acid import may impact on chemoresistance in breast cancer patients, we looked at the correlation between the co-expression of SLC3A2 and SLC7A5 and chemotherapy response using transcriptomic data from 3,104 breast cancer patients ^35^. We found that the two genes were highly expressed in chemoresistance (non-responder) tumours (**Figure 9E**) and the area under the ROC curve (AOC) was 0.628 (**Figure 9F**), classifying them as weak biomarkers with potential use in prediction of chemotherapy treatment ^35^.

**Figure 9.**
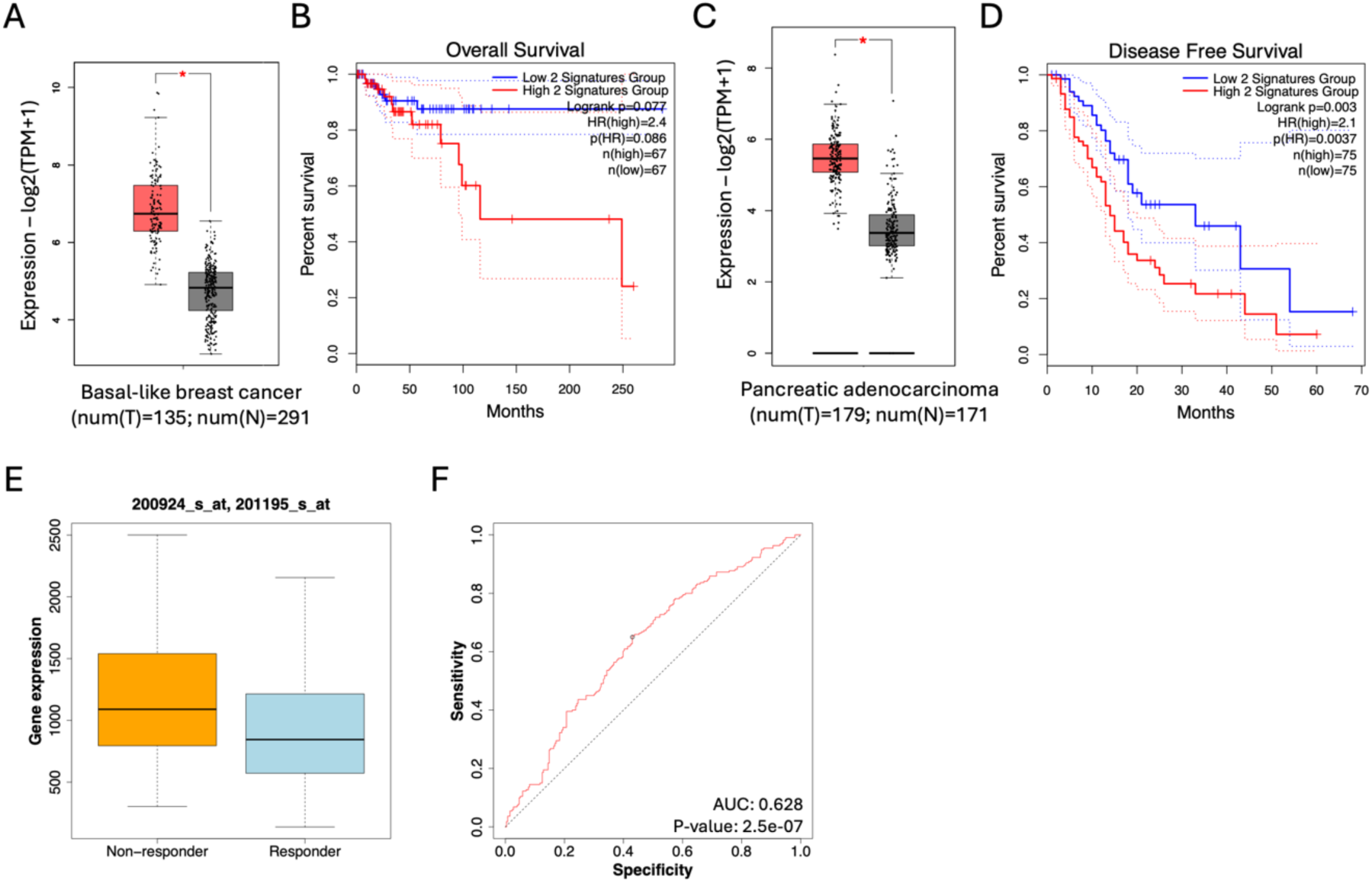
SLC3A2 and SLC7A5 co-expression was upregulated in breast and pancreatic tumours and correlated with poor prognosis and chemotherapy resistance. **(A,C)** RNA sequencing data from basal-Like breast tumours (n=135) and normal breast tissue (n=291) or pancreatic adenocarcinoma tumours (n=179) and normal pancreatic tissue (n=171) for the co-expression of SLC3A2 and SLC7A5. *p< 0.05. Data were obtained from gepia2.com. (B) Overall survival of basal-like breast cancer patients with high (red) or low (blue) co­expression of SLC3A2 and SLC7A5. Data were obtained from gepia2.com. (D) Disease free survival of pancreatic adenocarcinoma patients with high (red) or low (blue) co-expression of SLC3A2 and SLC7A5. Data were obtained from gepia2.com. (E,F) RNA sequencing data and ROC analysis for the co-expression of SLC3A2 and SLC7A5 from chemoresistant (Non-responder) and chemosensitive (Responder) breast tumours. (E) p = 1,4e-6, Mann-Whitney test. (D) AUG = 0.628; ROC p value = 2.5e-07. Data were generated in ROCplot.com.

## Discussion

Here, we showed that the ECM component collagen I supported the growth/survival of invasive breast cancer (MDA-MB-231 and MCF10CA1) and pancreatic cancer cells (PANC1 and SW1990) under Glc starvation. In contrast, non-invasive breast cancer (MCF10DCIS) and non-transformed mammary epithelial cells (MCF10A) showed no such growth advantage in the presence of collagen I. These findings suggest that the ability to utilise collagen I for survival under nutrient deprivation is a mechanism acquired during cancer progression and associated with invasiveness. We also observed a varying effect on cell behaviour; in PANC1, SW1990, and MCF10CA1 cells, collagen I promoted both cell division and survival, whereas in MDA-MB-231 cells, it supported survival, without promoting cell division. PANC1 cells are classified as Glucose Insensitive Cells (GIC) due to their strong antioxidant capacity, which confers high tolerance to reactive oxygen species (ROS) and enables them to maintain stable ATP levels. Conversely, MDA-MB-231 cells are considered Glucose Sensitive Cells (GSC) due to a lower tolerance to metabolic stress, characterised by a failure of ROS regulation and a more pronounced decrease in ATP levels^36^. Therefore, we hypothesise that the fundamental role of collagen I engagement via integrin receptors is to provide a potent survival signal that activates anti-apoptotic pathways, such as FAK/PI3K/Akt signalling ^37^. Cell proliferation is extremely energy-intensive and requires a high supply of ATP and building blocks ^38^. In this context, GICs like PANC1 cells are fully capable of supporting the increased energy demand and higher ROS production inherent to rapid growth, resulting in a successful increase in proliferation. Meanwhile, GSCs, such as MDA-MB-231 cells, are limited by their metabolic fragility, which prevents them from sustaining the intense energy expenditure required for rapid, active proliferation, thus resulting in an unchanged division rate.

We demonstrated that the mechanism through which collagen I supported cell growth/survival under Glc starvation was independent from ECM internalisation. Our recent work demonstrated that breast cancer cells relied on ECM uptake followed by lysosomal degradation to grow under amino acid starvation ^24^, suggesting that collagen I promotes cell growth/survival through different mechanisms depending on the type of nutrient limitation. Furthermore, our results contrast with studies showing that soluble albumin or collagen did not support pancreatic cancer cell growth under Glc deficiency ^39, 40^. However, it has been shown that α2β1 integrin has higher affinity for fibrillar collagen I, compared to soluble collagen^41^, suggesting that matrix collagen I might be required to trigger α2β1-dependent cell survival. Indeed, here we showed that inhibition of the major collagen I-binding integrin receptor, α2β1^42^, significantly reduced cell growth and invasion in both breast and pancreatic cancer cells under Glc starvation.

Mechanistically, we showed that collagen I promoted 2 separate pathways: the activation of mTORC1, a central sensor of nutrient availability ^43^, and the stimulation of essential amino acid import (**Figure 10**). Interestingly, prolonged Glc starvation reduced 4EBP1 phosphorylation, but not S6 in breast cancer cells. While Glc deprivation has been established to inhibit mTORC1 signalling ^27^, this response has mostly been characterised after short-term Glc limitation (30 minutes to 3 hours). Therefore, it is possible that the longer starvation used in this study might result in additional cellular adaptations, leading to the reactivation of S6 phosphorylation, but not 4EBP1. Consistent with this, S6K phosphorylation has been shown to be induced by Glc starvation in muscle cells ^44^. However, collagen I promoted S6 phosphorylation regardless of nutrient availability and was consistently reduced by α2β1 integrin pharmacological inhibition or siRNA-mediated knockdown. These results are in line with previous studies demonstrating that integrin-ECM binding activates the Akt/mTORC1/S6K/pS6 pathway ^45, 46^.

**Figure 10.**
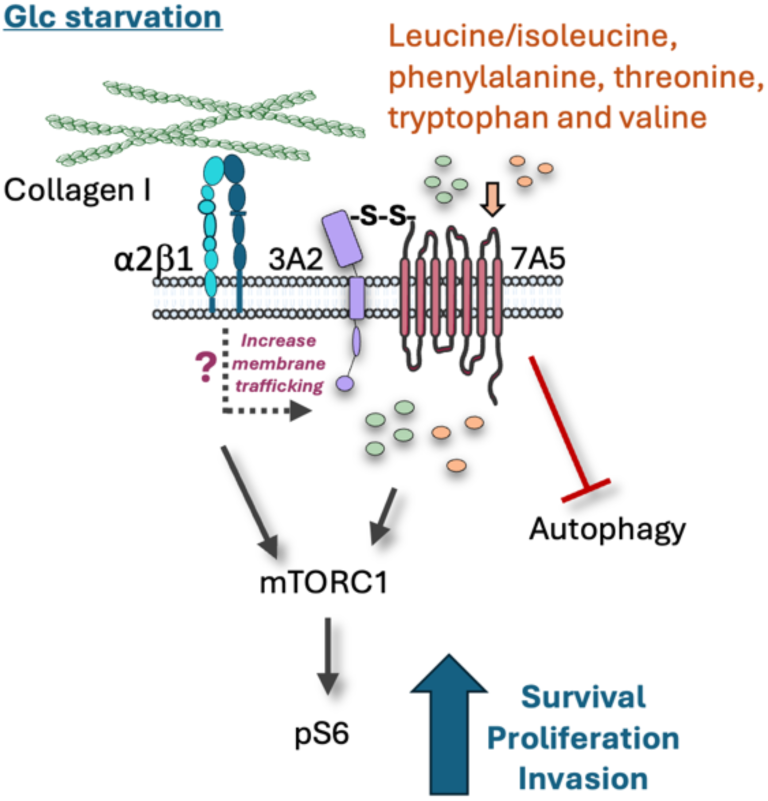
Working model. Under Glc starvation, collagen I promoted α2β1 integrin-dependent mTORCI activation in breast and pancreatic cancer cells. Collagen I also increased the membrane levels of the LAT1-4F2hc amino acid transporter heavy chain, SLC3A2 (3A2), and light chain, SLC7A5 (7A5), stimulating amino acid import. This resulted in mTORCI activation and autophagy inhibition, leading to increased survival, proliferation and invasion, in 2D and 3C contexts.

Glc starvation is a well-documented inducer of autophagy, a process of cellular self-digestion ^47^. While this held true for cells grown on plastic, we observed that cells cultured on collagen I exhibited reduced LC3 recruitment to autophagosomes under Glc starvation. Integrin activation has been shown to prevent autophagy via several pathways, including the PI3K-Akt/mTOR ^48^ and Src/FAK ^49^ pathways, which deactivate AMPK, a major autophagy regulator ^50^. However, this relationship is not always straightforward, as evidenced by studies where fibronectin secreted from skeletal muscle activated hepatic autophagy in an α5β1 integrin-dependent manner ^51^. Interestingly, our results show a more complex regulatory landscape. The α2β1 integrin inhibitor, BTT-3033, and the mTORC1 inhibitor, Rapamycin, did not significantly alter autophagy in cells seeded on collagen I under Glc starvation, suggesting that the observed reduction in autophagy in the presence of collagen I is not directly mediated by α2β1 integrin or mTORC1 activation. This implies that, while integrin signalling and mTORC1 are essential for collagen I-dependent growth/survival, the downstream suppression of autophagy may be mediated by an independent pathway. As a distinct collagen I receptor, discoidin domain receptor 2 (DDR2) offers a parallel signalling route to integrins, with evidence showing DDR2-mediated autophagy inhibition in adventitial fibroblasts ^52^. Thus, DDR2 could provide a redundant, non-integrin-dependent mechanism for guaranteeing the suppression of catabolic autophagy necessary for proliferation and survival.

The activation of mTORC1 has been shown to occur through several signalling pathways. One such mechanism is triggered by the presence of essential amino acids in the cells, particularly leucine ^43^. It is well established that amino acid-dependent activation of mTORC1 is primarily mediated by their cellular uptake by plasma membrane transporters. The LAT1-4F2hc complex (also known as SLC7A5-SLC3A2) is central to this mechanism ^34, 53, 54^. In our study, in both breast and pancreatic cancer cells under Glc starvation, collagen I resulted in increased intracellular essential amino acids, including isoleucine, leucine, phenylalanine, methionine, tryptophan, tyrosine, and valine, known substrates for the LAT1-4F2hc transporter ^33^. We demonstrated that collagen I increased the membrane localisation of both SLC3A2 and SLC7A5 in breast and pancreatic cancer cells under Glc deficiency, which provides a mechanistic explanation for the increased intracellular amino acid content observed. SLC3A2 is critical for the proper function and localisation of SLC7A5, acting as an ancillary protein that supports SLC7A5 traffic to the plasma membrane. The N-glycosylation of SLC3A2 is crucial for its own stability and successful transport, which in turn ensures the correct localisation of SLC7A5 ^55–57^. Moreover, the association with SLC3A2 is fundamental for the transport activity of SLC7A5, as SLC7A5 alone cannot efficiently transport amino acids ^57^. Consistently, our data show that knocking down both SLC3A2 and SLC7A5 has synergistic effects on intracellular amino acid levels. Additionally, downregulation of SLC3A2 decreased the phosphorylation of S6, an mTORC1 downstream target, and increased the autophagy marker LC3-II in both MDA-MB-231 and PANC1 cells. This is consistent with previous observations showing that LAT1 inhibition decreased S6 phosphorylation and promoted autophagy ^34, 58^.

SLC3A2 and integrin crosstalk has been previously reported. SLC3A2 is known to physically associate with integrins, particularly with the cytoplasmic domain of β1 integrin in mammalian cells and βPS in Drosophila, and this interaction drives adhesion signalling ^59, 60, 59, 61, 62^. Moreover, SLC3A2 has been shown to regulate integrin distribution in polarised cells ^63^, while it is currently unknown whether integrin activation controls SLC3A2 function/localisation. Further research is needed to investigate whether the activation of ECM-bound integrins could provide a binding domain on the cell membrane for SLC3A2, and consequently, SLC7A5. This potential interaction could offer a novel mechanism for how ECM signals regulate amino acid transport.

It remains unclear whether the α2β1 and the LAT1-4F2hc pathways are activated independently upon collagen I adhesion or are interlinked. Previous studies have shown that the phosphorylation of 4EBP1 downstream of mTORC1 transcriptionally regulates amino acid transporters by controlling the translation and stability of the transcription factor ATF4 ^64^. However, we did not observe any difference in the expression of SLC3A2 and SLC7A5 in the presence of collagen I compared to plastic, and α2 integrin knockdown did not alter SLC3A2 total protein level. Additionally, there was no difference in 4EBP1 activation between collagen I and plastic, suggesting that this mechanism might not be at play in our context. We can hypothesise that collagen I binding might affect LAT1-4F2hc membrane trafficking and localisation. Integrins have been shown to control protein membrane localisation, primarily by modulating the endocytosis, recycling, and exocytosis machinery. For instance, integrin engagement with the ECM regulates the surface density of Kir2.1 ion channel, which is preferentially internalised by endocytosis at focal adhesion sites ^65^. Furthermore, integrins act as mechanical sensors that trigger the rapid delivery of proteins, such as the Kv1.5 potassium channel, from Rab11-associated recycling endosome to the cell membrane ^66^. More broadly, integrin signalling controls the spatial and temporal turnover of membrane receptors, including integrins themselves, by regulating key trafficking components like Rab GTPases and phosphoinositides ^67^. We can therefore envisage a mechanistic framework linking integrin activation by collagen I to the control of amino acid transporter activity and localisation. Detailing the specific molecular targets within this pathway is now the critical next step and promises to uncover further novel therapeutic targets against matrix-driven diseases.

Evidence in the literature supports the role of both SLC3A2 and SLC7A5 in promoting tumour growth and survival, as their elevated expression enables cancer cells to thrive in a hostile tumour microenvironment. Upregulated expression of SLC3A2 in osteosarcoma and gliomas promotes tumour growth by activating the PI3K/Akt pathway ^68^ and is linked to enhanced malignancy and poor prognosis ^69^. Similarly, high expression of SLC7A5 is a common feature across several cancers ^70^. In bladder cancer, it is associated with poor prognosis and enhanced proliferation, migration, and invasion ^71^, while in pulmonary adenocarcinoma, it predicts patient prognosis ^72^. The inhibition of SLC7A5 has shown promise in suppressing cancer progression. In melanoma, knockdown or inhibition of SLC7A5 suppresses metastasis and proliferation by downregulating the mTOR signalling pathway ^73^. This is further supported by the observation that high SLC7A5 expression correlates with worse patient outcomes ^74^ and that its inhibition reduces global translation in cancer cells ^75^. The combined expression of SLC7A5 and SLC3A2 has also been identified as a potential predictor for response to endocrine therapy in oestrogen-receptor-positive breast cancer, where SLC7A5 expression was also correlated with genes related to proliferation and hypoxia ^76, 77^. In our study, knocking down SLC3A2 and SLC7A5 in invasive breast cancer cells inhibited 2D cell growth and 3D cell invasion. In PDAC cells, knocking down SLC7A5, but not SLC3A2, decreased growth in 2D, while the double knockdown of both genes decreased growth in 3D spheroids. It is important to consider that SLC3A2 functions as a chaperone for other amino acid antiporters, notably SLC7A11. The literature suggests that SLC7A11-mediated glutamate efflux depletes intracellular glutamate, thereby limiting the cell’s ability to utilise glutamine-derived carbon (via α-ketoglutarate) to maintain the tricarboxylic acid cycle in Glc-depleted conditions ^78^. This suggests that in the 2D PANC1 model, SLC3A2 knockdown may be affecting a compensatory metabolic mechanism mediated by SLC7A11 or other SLC3A2-dependent transporters, which ultimately offsets the growth reduction expected from decreased amino acid uptake.

In conclusion, our study demonstrated that collagen I partially rescued breast and pancreatic cancer cell growth under Glc deficient conditions, via a combination of two pathways. Firstly, α2β1 integrin was required to sustain mTORC1 activity, thereby promoting an anabolic state. Secondly and concurrently, the cells exhibited higher levels of the LAT1-4F2hc (SLC7A5-SLC3A2) transporter on the plasma membrane, resulting in increased amino acid uptake in the presence of collagen I under Glc starvation. This resulted in reduced autophagy and sustained mTORC1 activity. Therefore, targeting LAT1-4F2hc-dependent amino acid transport could represent a novel therapeutic strategy to prevent the growth and progression of highly fibrotic and nutrient deprived tumours.

## Methods

### Reagents

Primary antibodies for Phospho-S6 Ribosomal Protein ser235/236 and LC3A/B were from Cell Signalling; α2 integrin (CD49b)-FITC conjugated and β1 integrin (CD29)-Alexa Fluor 488 conjugated from BioLegend; α2 integrin for Western blotting from BD-bioscience; SLC3A2 (CD98) and SLC7A5 (LAT1) from Proteintech and GAPDH from SANTA CRUZ Biotechnology. Secondary antibodies Alexa-Fluor 594 anti-Rabbit IgG and Alexa-Fluor 488 anti-Rabbit IgG were from Cell Signalling; IRDye® 800CW (anti-mouse IgG), IRDye® 680CW (anti-rabbit IgG) and DR were from LI-COR. Alexa Fluor TM 488, 555 and 647 Phalloidin, Click-iT EdU Imaging Kits, Propidium iodide (PI), NHS-Fluorescein and Hoechst 33342 were from Invitrogen. Rat tail Collagen I (high concentration) was from Corning. All media and dialyzed FBS were from Gibco. E64d (Aloxistatin) was from AdooQ Bioscience. BTT3033 and VPS34-IN were from Cambridge Bioscience. D-phenylalanine was from Santa Cruz Biotechnology. Rapamycin was from Fluorochem.

### Cell culture

MDA-MB-231, PANC1 and SW1990 cells were mainteined in high glucose Dulbecco’s Modified Eagle’s Medium (DMEM) supplemented with 10% foetal bovine serum (FBS) and 1% penicillin/streptomycin (PS). MCF10A, MCF10A-DCIS and MCF10CA1 cell lines, a gift from Professor Giorgio Scita (IFOM, Milan), were cultured in DMEM/F12. MCF10A-DCIS cells were supplemented with 5% Horse serum (HS), 20 ng/ml epidermal growth factor (EGF), and 1% PS. MCF10A and MCF10CA1 cells were grown with 5% HS, 20 ng/ml EGF, 0.2 mg/ml hydrocortisone, 10 μg/ml insulin, and 1% PS. All cell lines were incubated at 37°C with 5% CO_2_ and passaged every 3 to 4 days.

To culture E0771 mouse triple negative breast organoids, 24-well plates were coated with a 10µl drop of 10.3 mg/ml Matrigel, polymerised for 10 minutes at 37°C to make a dome. Organoids were diluted in 10.3 mg/ml Matrigel and 10µl of Matrigel containing organoids were seeded on top of the dome and incubated upside down at 37°C for 30 minutes. Then 1ml of advanced DMEM/ F12 supplemented with 1% PS, GlutaMAX (1% vol/vol), HEPES (1% vol/vol), hEGF (5 ng/ml), hydrocortisone (0.3 μg/ml) and 10 µM Y-27632 were added. Organoids were incubated at 37°C with 5% CO_2_ and passaged every 7 days.

### ECM preparation

Collagen I was diluted in ice-cold PBS to final concentrations of 2 mg/ml and incubated at 37°C with 5% CO_2_ for 3 hours to polymerise.

### ECM cross-linking

Polymerised collagen I was treated with 10% glutaraldehyde for 30 minutes at room temperature (RT). Following two PBS washes, the glutaraldehyde was quenched with 1M glycine for 20 minutes at RT, followed by two additional PBS washes. Cross-linked collagen I was then stored overnight at 37°C with 5% CO_2_ in PBS.

### Starvation conditions

Glucose- and pyruvate-free media DMEM was supplemented with 10% dialyzed FBS (DFBS) for MDA-MB-231, PANC1 and SW1990 cells. Glucose- and pyruvate-free media DMEM/F12 was supplemented with 5% HS, 10 μg/mL insulin and 20 ng/mL EGF for MCF10A cells; 2.5% HS, 1% PS, 20 ng/ml EGF, 10 μg/ml insulin, 0.2 mg/ml hydrocortisone for MCF10CA1; 5% HS and 20 ng/mL EGF for MCF10A-DCIS cells.

### Proliferation assay

96-well plates were coated with collagen I at a concentration of 2 mg/ml, 15 µl per well. Three wells were designated as technical replicates for each experimental condition.

10^3^ MDA-MB-231, MCF10A, MCF10A-DCIS, PANC1 cells/well, 2×10^3^ SW1990 cells/well and 400 MCF10CA1 cells/well were seeded in their respective complete growth media. After a 5-hour incubation at 37°C in a 5% CO_2_ atmosphere, the media was replaced with 200 µl of glucose starvation media or complete media with 10% DFBS. Where indicated, the starvation media also contained 5, 10 or 15 µM BTT-3033 or DMSO as vehicle control. Inhibitors were replenished every two days for a total duration of four days. Cells were subsequently fixed by the addition of 4% paraformaldehyde (PFA) for 15 minutes RT, followed by two washes with PBS. Nuclear staining was performed using either DR or Hoechst 33342. 5 mM DR in PBS was applied to the cells for 1 hour at RT with gentle rocking. Cells were then washed twice with PBS for 30 minutes to minimize background fluorescence before imaging. DR fluorescence was detected using a Licor Odyssey Sa instrument via the 700 nm channel with 200 mm resolution. Signal intensity (calculated as total intensity minus total background) for each well was quantified using Image Studio Lite software. Alternatively, cells were fixed directly with 4% PFA containing 10 µg/ml Hoechst 33342 for 15 minutes, followed by two PBS washes. Images were acquired with an ImageXpress Micro using a 2x objective, ensuring full well coverage of each well. Image analysis was conducted using MetaXpress and Costume Module Editor (CME) software at the Sheffield RNAi Screening Facility (SRSF)

### EdU incorporation assay

96-well plates were coated with 2 mg/ml collagen I. 10^3^ MDA-MB-231 cells/well, 2×10^3^ PANC1 and SW1990 cells/well and 400 MCF10CA1cells/well were seeded in complete growth media. After a 5-hour incubation at 37°C with 5% CO_2_, the complete growth media was replaced with 200 µl of glucose starvation media. At designated time points (day six post-starvation for MDA-MB-231 cells, day five for PANC1, day seven for SW1990 cells and day four for MCF10CA1), cells were incubated with 5 mM EdU for 1-2 days, at 37°C with 5% CO_2_. Cells were then fixed using 4% PFA containing 10 µg/mL Hoechst 33342 for 15 minutes at RT, and subsequently permeabilized with 0.25% Triton X-100 for 5 minutes. EdU detection was achieved by incubating cells with the Click-iT EdU Alexa Fluor 555 detection cocktail (Invitrogen) or EdU Cell Proliferation Image kit (Antibodies.com) for 30 minutes at RT with gentle rocking. Following two PBS washes, cells were stored in PBS for imaging. Images were acquired using an ImageXpress Micro system with a 2x objective, and subsequent quantification was performed using MetaXpress and Costume Module Editor (CME) software.

### Propidium iodide (PI) live-cell imaging

MDA-MB-231, SW1990 and PANC1 cells were seeded in 96-well plates coated with 2 mg/ml collagen I in complete growth media. Following a 5-hour incubation period at 37°C with 5% CO_2_, the complete growth media was replaced with glucose starvation media. At day 4, 5, and 6 or 7 post-starvation, cells were co-incubated with 1 µg/ml Propidium Iodide (PI) and 5 µg/mL Hoechst 33342 for 30 minutes at 37°C with 5% CO_2_. Cells were then washed once with PBS and maintained in phenol red-free media for imaging. Images were acquired live using an ImageXpress Micro system with a 10x objective, and subsequent quantification was performed with MetaXpress and CME software.

### Cleaved Caspase 3/7 staining

MCF10CA1 cells were seeded 96-well plates coated with 2 mg/ml collagen I in complete growth media. Following a 5-hour incubation at 37°C with 5% CO_2_, the complete growth media was replaced with glucose starvation media. At day 6 post-starvation media was changed to PBS containing 5 μM Cell Event Caspase-3/7 Green Detection Reagent for 1 hour and 30 minutes. Cells were fixed and stained with Hoechst 33342. Images were collected using an Image Xpress micro system with a 10x objective and quantified with MetaXpress and CME software.

### Immunofluorescence

For immunofluorescence staining of α2 integrin, β1 integrin and pS6 1×10^4^ cells were seeded in 96-well plates. For SLC3A2 and SLC7A5 staining, 2×10^5^ cells were seeded in 3.5 cm glass-bottom dishes. Cells were fixed with 4% PFA for 15 minutes, followed by two washes with PBS. For surface staining of α2 integrin, β1 integrin, SLC3A2, and SLC7A5, cells were not permeabilized. Instead, they were directly incubated with 3% bovine serum albumin (BSA) in PBS for 1 hour at RT for blocking. For pS6 staining, cells were permeabilized with 0.25% Triton X-100 for 5 minutes at RT, followed by washes and subsequent blocking with 3% BSA in PBS for 1 hour at RT. Following blocking, cells were incubated with primary antibodies (1:200 dilution) for 2 hours at RT, followed by two PBS washes. Cells were then incubated with an Alexa Fluor 488 or Alexa Fluor 555 anti-rabbit secondary antibody (1:1000) and Phalloidin Alexa Fluor 555 or Alexa Fluor 647 (1:1000) for 1 hour at RT, followed by two final PBS washes. Cells were stained with 5 µg/mL Hoechst 33342 and kept in PBS for imaging. α2 and β1 integrin images were collected by an ImageXpress Micro system with a 10x objective, while SLC3A2, SLC7A5 and pS6 images were collected by a Zeiss LSM980 Airyscan 2 System with a 40x 1.2 NA water (SLC3A2 and SLC7A5) or a 20x 0.8 NA (pS6) objective.

### ECM uptake

3.5 cm glass-bottomed dishes were coated with 2 mg/ml collagen I and incubated for 3 hours at 37°C with 5% CO_2_ for polymerization. Collagen I was then labelled with 10 µg/ml Fluorescein succinimidyl ester (NHS) for 1 hour at RT on a gentle rocker. Following labelling, 1×10^5^ MDA-MB-231 cells/dish were seeded in complete growth media. After a 5-hour incubation at 37°C with 5% CO_2_, the complete growth media was replaced with 1 ml of glucose starvation media. Cells were treated with either 20 μM E64d or DMSO. Cells were fixed on day three by adding 4% PFA for 15 minutes at RT. Fixed cells were permeabilised with 0.25% Triton X-100 for 5 minutes and then incubated with Phalloidin Alexa Fluor 555 (1:400 in PBS) for 10 minutes to visualise actin. Dishes were mounted with 2-3 drops of Vectashield mounting medium containing DAPI and sealed with parafilm, then stored at 4°C. Cells were visualised using a Nikon A1 confocal microscope equipped with a 60x 1.4 NA oil immersion objective. Collagen I uptake index was quantify with Fiji/ImageJ ^79^ as in Nazemi et al., 2024 ^24^.

### Western blotting

MDA-MB-231, MCF10CA1 and PANC1 cells were lysed with lysis buffer (50 mM Tris (pH 7) and 1% SDS) and transferred to Qiashredder columns (Qiagen) to remove the DNA. Lysates were diluted 4:1 in NuPAGE-LDC sample buffer and run on 4% to 15% Mini Protean TGX gels in SDS-PAGE running buffer (3.03 g Tris base, 14.41 g glycine, and 1 g SDS) at 100 V for 1 hour and 15 minutes. Proteins were transferred to Immobilon-FL PVDF membrane (MERCK) in transfer buffer (10% Towbin buffer, 20% methanol) at 100 volt for 75 minutes. Membranes were blocked in TBS-T with 5% w/v skimmed milk or 5% w/v BSA. Then, membranes were incubated with 1:1,000 GAPDH or 1:1000 α-tubulin together with primary antibodies for LC3, pS6, S6, α2 integrin and SLC3A2 1:1000 in TBS-T with 5% w/v skimmed milk or 5% w/v BSA overnight at 4°C. Membranes were incubated with secondary antibodies, IRDye 800CW anti-mouse IgG for α2 integrin, GAPDH and α-tubulin (1:30,000) and IRDye 680CW anti-rabbit IgG for LC3, pS6, S6 and SLC3A2 (1:20,000) in TBS-T with 0.01% SDS, for 1 hour at RT on the rocker. Membranes were washed 2 times in TBS-T for 15 minutes on the rocker at RT. Images were taken by a Licor Odyssey Sa system. Band intensity was quantified with Image Studio Lite software.

### Non-targeted metabolite profiling

MDA-MB-231, MCF10CA1, SW1990 and PANC1 cells were cultured on collagen I-coated 6-well plates in complete growth media. Following a 5-hour incubation, the media was changed to glucose starvation media for up to 4 days. Metabolite extraction involved washing cells with ice-cold PBS, followed by incubation with a cold 5 MeOH: 3 AcN: 2 H_2_O solution. After centrifugation, samples were analysed by electrospray ionization mass spectrometry (Waters G2 Synapt) at the Sheffield Faculty of Science Biological Mass Spectrometry Facility (biOMICS), with data acquired in both positive and negative ion modes. Three technical replicates per sample were run, and only peaks consistently present across all replicates were included. Data was binned (0.2 amu m/z), and putative metabolite were identified using the HumanCyc database. Data analysis was performed using Perseus software (v1.5.6.0) and used Student t test (SO = 0.1, false discovery rate (FDR) = 0.05 and p<0.05), and metabolic pathway analysis was conducted with MetaboAnalyst 5.0 (p<0.05).

### Targeted

Metabolomics. Metabolites extraction was performed as above. A mass spectrometer Waters Synapt G2-Si coupled to Waters Acquity UPLC was used to separate alanine, asparagine, aspartic acid, arginine, cysteine, glutamic acid, glycine, glutamine, histidine, isoleucine, lysine, leucine, phenylalanine, methionine, serine, proline, tryptophan, threonine, tyrosine and valine with separation was carried out on a Waters BEH C18 column (50 x 2.1 mm) at 40°C, with an injection volume of 5 μl. The mobile phases consisted of 0.1% formic acid (A) and 0.1% acetonitrile (B). The flow rate was 0.4 ml/min, and the gradient program was as follows: 1–35% B (0–3 min), 35-99% B (3–6 min), 99% B (6-6.9 min), 99-1% B (6.9-7 min), and 1% B (7-8 min). The samples were run in positive and negative modes. To identify the targeted metabolites, the retention time and the mass of the compound were matched with the standards. A minimum of three independent biological replicates were performed, with one well of the 6-well plate dedicated to each experimental condition per replicate.

### siRNA Transfection

For cell proliferation experiments, Dharmacon ON-TARGETplus siRNA smart pools (150 nM, 20 µl per well) were mixed with Dharmafect 1 (DF1, 0.24 µl in 19.76 µl DMEM, 20 µl per well) and incubated in a 96-well plate for 30 minutes at RT. Following this, MDA-MB-231 and PANC1 cells were seeded in 60 µl of 10% FBS DMEM (antibiotic-free), while MCF10CA1 cells were seeded in 5% HS DMEM/F12, establishing a 30 nM final siRNA concentration. Cells were maintained overnight at 37°C and 5% CO_2_. Media was then changed to 200 µl of either complete (with DFBS) or glucose starvation media for up to four days. Cells were fixed, stained with 10 µg/ml Hoechst 33342, and imaged using ImageXpress Micro. Images were analysed with MetaXpress and CME software.

For western blotting, 3D spheroids and metabolite profiling, in a 6-well plate, 10 µl of 5 µM siRNA was mixed with 190 µl of Opti-MEM per well. Separately, 2 µl of DF1 was combined with 198 µl of Opti-MEM and incubated for 5 minutes at RT. The Opti-MEM/DF1 mixture (200 µl) was then added to the siRNA solution and incubated on a rocker for 20 minutes. Finally, cells in 1.6 ml of media were added to each well. After 24 hours of incubation at 37°C and 5% CO_2_, the media was replaced with 3 ml of glucose starvation media.

**Table 1.** siRNA smart pools used in this study.

| Type | Target | Ref No. |
| --- | --- | --- |
| si-GENOME siRNA smartpool | Human SLC7A5 | M-004953-01-0005 |
| **ON-TARGETplus siRNA smartpool** | Human SLC3A2 | L-003542-00-0005 |
| **ON-TARGETplus siRNA smartpool** | Human ITGA2 | L-004566-00-0005 |
| **ON-TARGETplus siRNA smartpool** | Human ITGB1 | L-004506-00-0005 |
| **ON-TARGETplus siRNA** | Non-targeting siRNA #4 | D-001810-04-20 |

### RT-QPCR

mRNA was extracted from MDA-MB-231 and PANC1 cells using the Qiagen RNeasy® Mini kit, following manufacturer’s instructions. Cells grown on plastic were trypsinised, while those on collagen I were scraped; both were then pelleted, washed, and snap-frozen. Lysis in Buffer RLT and purification through spin columns, followed by washes, yielded purified mRNA. mRNA was then used for cDNA synthesis with the High-Capacity cDNA Reverse Transcription Kit (Fisher), where 1 μg of mRNA per sample was reverse transcribed in a 20 μl reaction containing RT buffer, dNTP mix, random primers, and MultiScribe Reverse Transcriptase. A negative reverse transcriptase control was included. The synthesized cDNA was stored at −80°C. Finally, qPCR analysis involved preparing a master mix with QuantiNova SYBR® Green PCR Kit and QuantiTect® Primer Assay. 7 μl of this mix was combined with 3 μl of 1:100 diluted cDNA (5 ng/μl final concentration) in a 384-well plate. -RT and blank water controls were included for all target genes. Samples were run on a Quantstudio 12K flex real-time PCR system for 40 cycles, with a melting curve analysis to confirm product specificity. Target gene expression was calculated using the 2−ΔCt method, normalizing to GAPDH as the housekeeping gene.

**Table 2.** RT-QPCR primers used in this study.

| **Primer** | **Assay name** | **Cat No.** |
| --- | --- | --- |
| **GAPDH** | Hs_GAPDH_1_SG | QT00052248 |
| **SLC3A2** | Hs_SLC3A2_1_SG | QT00085897 |
| **SLC7A5** | Hs_SLC7A5_1_SG | QT00089145 |
| **SLC7A6** | Hs_SLC7A6_1_SG | QT00002674 |
| **SLC7A8** | Hs_SLC7A8_1_SG | QT00006384 |

### Stable cell line generation

MDA-MB-231 cells were plated in 6-well plates and allowed to grow to high confluency. On day 2, a Lipofectamine 2000-DNA complex was prepared by combining diluted Lipofectamine (5 μl LF in 250 μl Opti-MEM per well) with the 2.5 μg pEGFP-C1+mRFP LC3 plasmid in 250 μl Opti-MEM, incubated 20 minutes, and then added to cells in antibiotic-free DMEM. After overnight incubation, the transfection solution was replaced with complete media. Cells were grown in the presence of 1000 μg/ml G418 2-3 days post-transfection, when cells reached 50-70% confluency. GFP- and RPF-positive single cell clones were sorted using a Sony MA900 cell sorter, selecting for cells with high intensity for both RFP and GFP.

### 3D Spheroid Generation

MDA-MB-231, stably expressing GFP as described in Nazemi et al., 2024 ^24^, and PANC1 cell 3D spheroids were generated via the hanging drop method. Each 20 μl drop contained 500 cells suspended in a mixture of 4.8 mg/ml methylcellulose and 20 μg/ml soluble collagen I. After 48 hours, spheroids were embedded in a matrix mix composed of 3 mg/ml collagen I and 3 mg/ml Geltrex. To avoid spheroids sinking to the bottom of the plate, the plate was slowly turned upside-down and incubated for 30 minutes at 37°C. Then 1 ml media were added to each well.

For drug treatment studies, the media was changed to glucose starvation media 24 hours post-embedding, and spheroids were subsequently treated with either 10 μM BTT-3033 or DMSO control. In siRNA knockdown experiments, cells were transfected for 24 hours prior to spheroid formation and embedding in the collagen I and Geltrex matrix.

MDA-MB-231 spheroids were imaged using a Nikon A1 confocal microscope with CFI Plan Fluor 10x (NA 0.3) objective. For drug treatment assays, imaging occurred from day 1 to day 3 post-embedding. For siRNA knockdown experiments, imaging was performed on day 1 and day 4 post-embedding. The invasion area was quantified by subtracting the spheroid’s core area from its total area.

PANC1 spheroids were imaged every two days up to day 6 post-starvation by an Olympus E450 microscope with 10x objective, with drug treatments being refreshed every two days.

### Mouse tumour organoid invasion

E0771 mouse triple negative breast tumour organoids were grown in 3 mg/ml collagen I and 3 mg/ml Gelterx (1:1). 24-well plates were coated with a 10 µl drop of 10.3 mg/ml Matrigel, polymerised for 10 minutes at 37°C to make a dome. 10 µl of organoids in Geltrex-collagen I mixture were added on top of the dome and incubated upside down at 37°C for 30 minutes. Then 1 ml of advanced DMEM/F12 supplemented with 1% PS, GlutaMAX (1% vol/vol), HEPES (1% vol/vol), hEGF (5 ng/ml), hydrocortisone (0.3 μg/ml) and 10 µM Y-27632 was added and organoids were incubated at 37°C with 5% CO_2._ After one day, the media was changed to glucose starvation media. Cells were treated with DMSO (Ctrl), 2.5 or 5 µM BTT-3033 or 50 mM D-phenylalanine. Cells were imaged live every day up to three days with an Olympus E450 microscope with 4x and 10x objectives. Size of organoids were measured by quantifying the core area of each organoid. Invasion length was quantified by measuring the length of invasive branches. The resulting data points reflect individual measurements, with each point in the graph representative of a single protrusion’s length.

### Expression, survival and ROC analysis

RNA sequencing data from basal-like breast cancer, pancreatic adenocarcinoma, normal breast and pancreatic tissue and survival analysis were performed in gepia2 (http://gepia2.cancer-pku.cn/#index), using TCGA/GTEx data. The link between gene expression and response to therapy (including taxane, anthracycline, ixabepilone, CMF, FAC and FEC) were analysed in ROC plotter (https://www.rocplot.com/), using transcriptome-level data of breast cancer patients ^35^. Breast cancer datasets were identified in GEO (https://www.ncbi.nlm.nih.gov/gds), using the platform IDs “GPL96”, “GPL570”, and “GPL571”. Jetset probes were used (https://services.healthtech.dtu.dk/services/jetset/).

### Statistical analysis

Graphs were generated using GraphPad Prism software (versions 10). To compare two datasets, a Mann-Whitney U test was employed. For more than two datasets with a single independent variable, one-way ANOVA (Kruskal–Wallis, Dunn’s multiple comparisons test) was applied. When two independent variables were present, two-way ANOVA (Tukey’s multiple comparisons test) was utilized. Metabolomics data were analysed in Perseus software.

## Acknowledgements

Imaging work was performed at the Wolfson Light Microscopy Facility, University of Sheffield (funded by the Wellcome Trust, grant WT093134AIA), using a Nikon A1 confocal microscope or a Zeiss LSM980 Airyscan 2 System. High-throughput imaging was performed in the RNAi facility at the University of Sheffield. Metabolomics analyses were performed in the biOMICS facility at the University of Sheffield. Fluorescence-Activated Cell Sorting (FACS) was carried out with the expert assistance of Dr Paul Gokhale at the Centre for Stem Cell Biology, University of Sheffield. qPCR analysis was performed in collaboration with the Tsakiridis lab at the University of Sheffield. We would like to thank Dr Eric Vancauwenberghe for extraction and generation of E0771 mouse triple negative breast tumour organoids, Prof Jasson King for sharing MDA-MB-231-LC3B-GFP cells, Dr Mark Collins for sharing the GFP-RFP-LC3 construct and Dr Montserrat Llanses Martinez for the generation of MDA-MB-231-GFP cells.

## Authors’ contributions

Conceptualisation: ER, MN; Data curation: MN; Formal analysis: MN, BY, IO, HW; Funding acquisition: ER; Investigation: MN, BY, IO, HW; Methodology: MN, BY, IO, HW; Project administration: ER; Supervision: ER; Writing, review and editing: MN, ER.

## Supplementary figures

**Figure S1.**
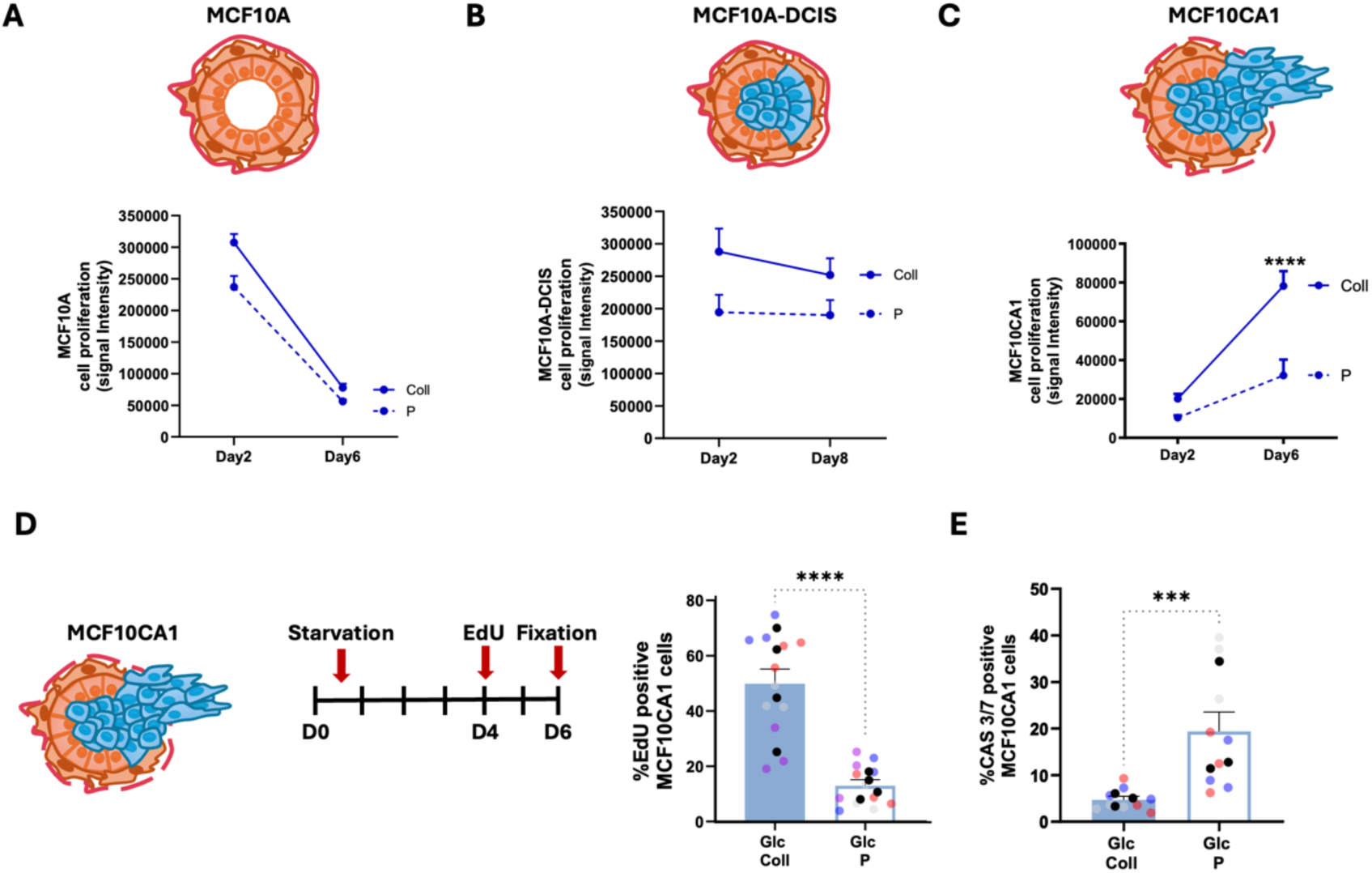
Collagen I supported cell growth of invasive breast cancer cells, but not non-invasive or non-transformed mammary epithelial cells. **(A-C)** MCF10A, MCF10A-DCIS and MCF10CA1 cells were seeded on plastic (P) or 2 mg/ml collagen I (Coll) for 6 or 8 days under Glc starvation, fixed, stained with DR, and imaged with a Licor Odyssey Sa system. Signal intensity was calculated by Image Studio Lite software. Data are presented as mean ± SEM, N=3 independent experiments, ****p < 0.0001 two-way ANOVA, Tukey’s multiple comparisons test. (D) MCF10CA1 cells were seeded on 2 mg/ml collagen I (Coll) or plastic (P) under Glc starvation. Cells were incubated with EdU at day 4 post starvation, fixed and stained with Hoechst 33342 and Click iT EdU imaging kit at day 6. **(E)** MCF10CA1 were seeded on plastic (P) or 2 mg/ml collagen I (Coll) for 6 days. Cells were fixed, stained for cleaved Cas3/7 and Hoechst 33342. Images were collected by ImageXpress micro and analysed by MetaXpress software. Data are presented as mean ± SEM, N≥3 independent experiments (the black dots represent the mean of individual experiments), ***p < 0.001, ****p < 0.0001 Mann-Whitney test.

**Figure S2.**
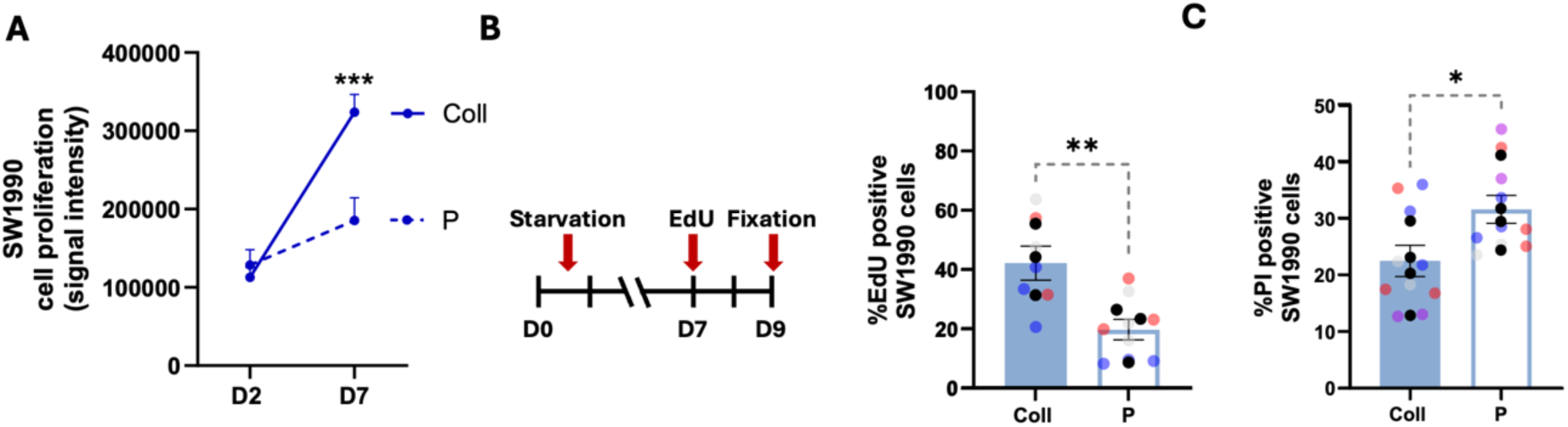
Collagen I supported cell growth and prevented cell death in SW1990 pancreatic cancer cells. **(A)** SW1990 cells were seeded on 2 mg/ml collagen I (Coll) or plastic (P) for 7 days under Glc starvation, fixed, stained with DR and imaged with a Licor Odyssey Sa system. Signal intensity was calculated by Image Studio Lite software. Data are presented as mean ± SEM, N=3 independent experiments, ***p < 0.001 two-way ANOVA, Tukey’s multiple comparisons test. (B) SW1990 cells were seeded on 2 mg/ml collagen I (Coll) or plastic (P) under Glc starvation, cells were incubated with EdU at day 7 post starvation, fixed and stained with Hoechst 33342 and Click iT EdU imaging kit at day 9. (C) SW1990 cells were seeded as in B and treated with PI at day 7. Images were collected by ImageXpress micro and analysed by MetaXpress software. Data are presented as mean ± SEM, N≥3 independent experiments (the black dots represent the mean of individual experiments), *p < 0.05, **p < 0.01, Mann-Whitney test.

**Figure S3.**
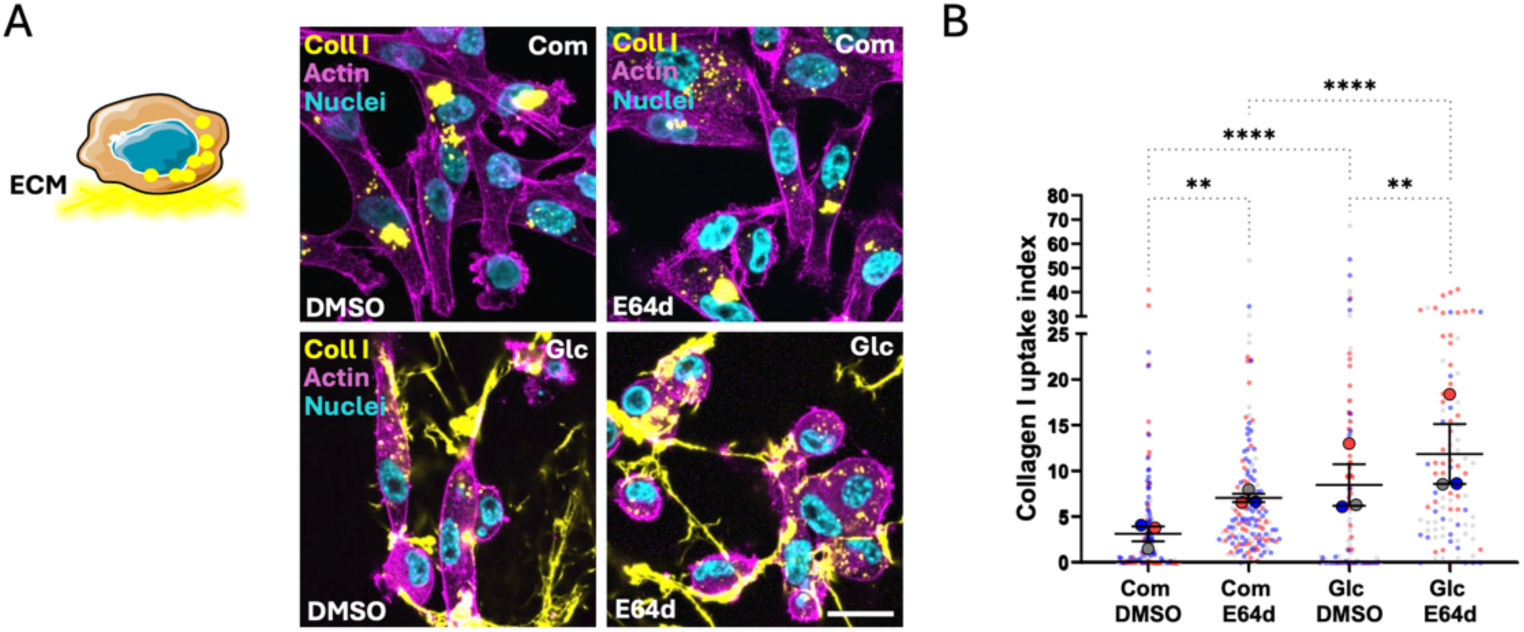
Glucose starvation promoted collagen I internalisation. **(A)** MDA-MB-231 cells were seeded on 2 mg/ml NHS-Fluorescein labelled collagen I (yellow) under complete (Com) or Glc starvation media in the presence of DMSO or 20 µM E64d for 3 days. Cells were fixed and stained for actin (magenta) and nuclei (cyan). Images were collected by Nikon Confocal A1 microscope. Bar, 20 µm. (B) Collagen I uptake index was measured with Fiji/lmageJ. Data are presented as mean ± SEM, N=3 independent experiments (the bigger dots represent the mean of individual experiments), **p < 0.01, ****p < 0.0001. One-way ANOVA Kruskal-Wallis, Dunn’s multiple comparisons test.

**Figure S4.**
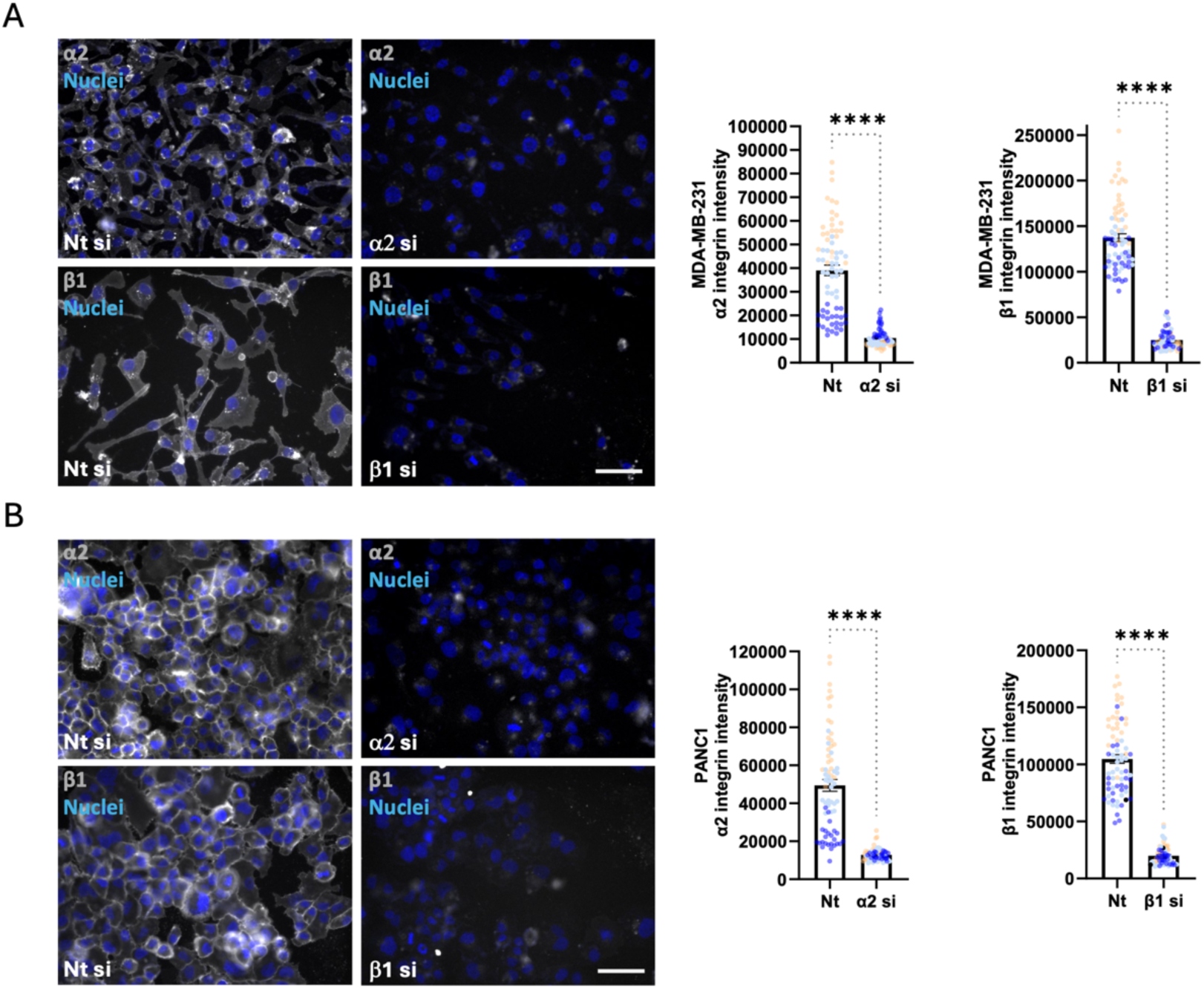
α2β1 integrin-targeting siRNAs significantly reduced protein expression. **(A)** MDA-MB-231 and **(B)** PANC1 cells were transfected with a non-targeting control siRNA (Nt si), an siRNA targeting α2 integrin (α2 si) or an siRNA targeting β1 integrin (β1 si) for 4 days. Cells were fixed and stained with Hoechst 33342 (blue) and α2 integrin or β1 integrin (grey). Images were collected by ImageXpress micro. Bar, 50 µm. Integrin signal intensity was quantified by MetaXpress software. Data are presented as mean ± SEM, N=3 independent experiments. p< 0.0001 Mann-Whitney test.

**Figure S5.**
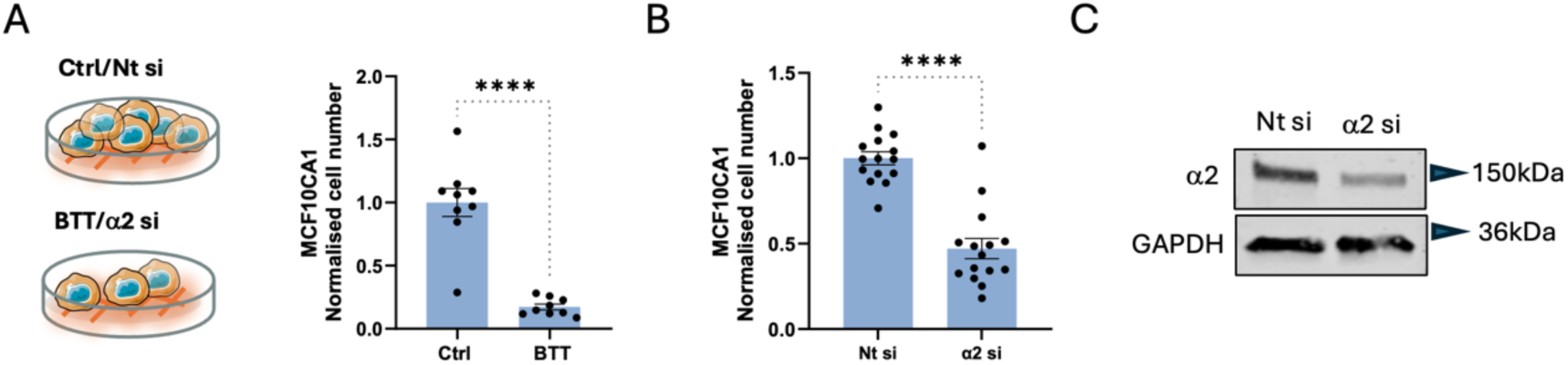
α2 integrin was required for MCF10CA1 cell growth. **(A)** MCF10CA1 cells were seeded on 2 **mg/ml** collagen I and grown under glucose starvation in the presence of 15 µM BTT-3033 (BTT) or DMSO (Ctrl) control for 4 days. (B) MCF10CA1 cells were transfected with a non-targeting control siRNA (Nt si) or an siRNA targeting α2 integrin (α2 si) and grown on 2 mg/ml collagen I under glucose starvation for 4 days. Cells were fixed and stained with Hoechst 33342. Innages were collected by ImageXpress micro. Data are presented as mean ± SEM, N=3 independent experiments. p < 0.0001 Mann-Whitney test. **(C)** MCF10CA1 cells were transfected as in B, lysed and the levels of α2 integrin and GAPDH were measured by Western blotting.

**Figure S6.**
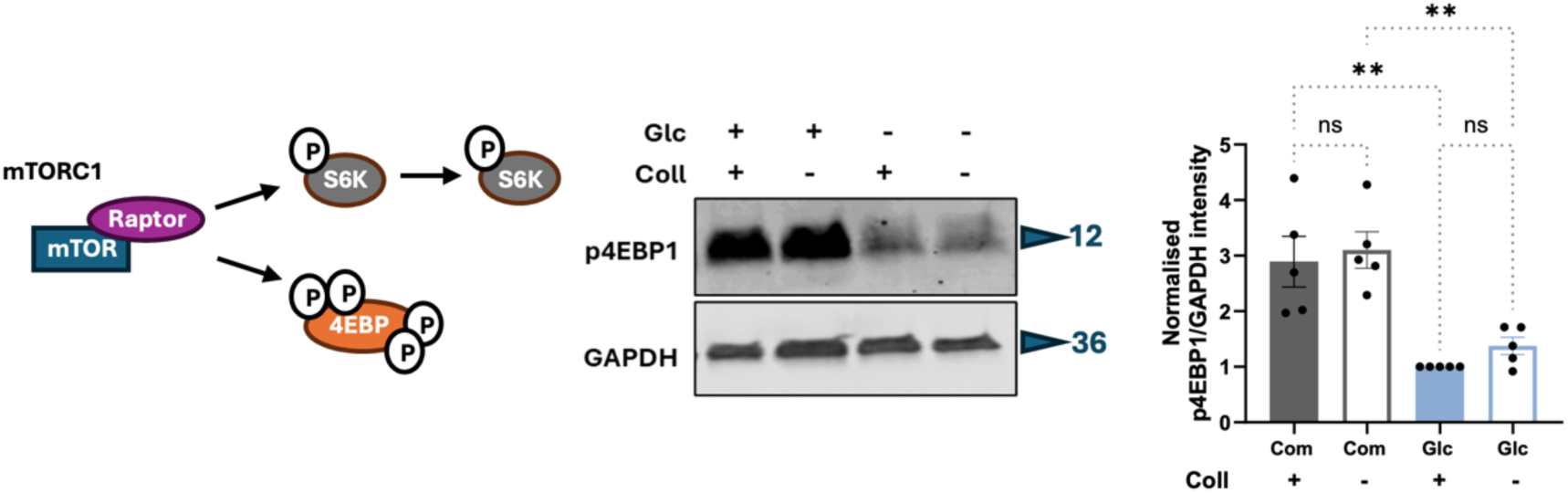
Collagen I did not affect the phosphorylation of the mTORCI target 4EBP1. MDA-MB-231 cells were seeded on 2 mg/ml collagen I (+) or plastic (-) under complete or Glc starvation media for 1 day. Lysates were collected and the levels of phosphorylated 4EBP1 (p4EBPI) and GAPDH were measured by Western blotting. Data are presented as mean ± SEM, N=5 independent experiments, **p < 0.01 Kruskal-Wallis, Dunn’s multiple comparisons test.

**Figure S7.**
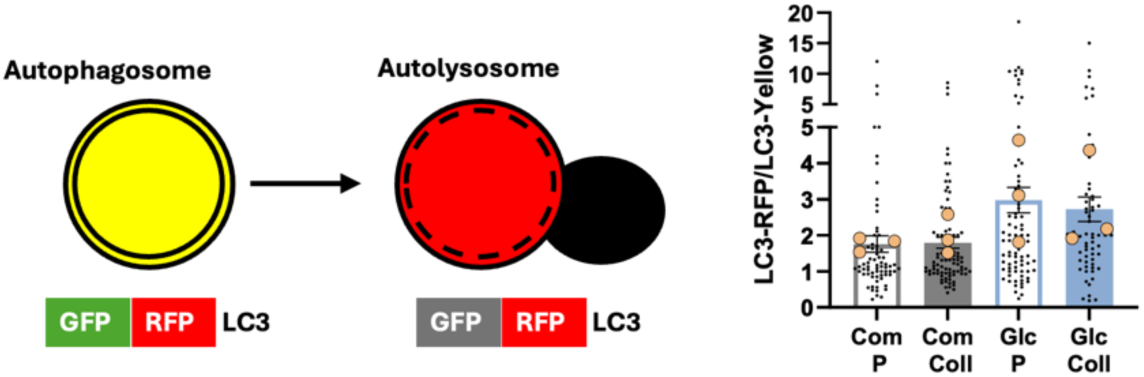
Collagen I did not affect autophagosome maturation. MDA-MB-231 cells stably expressing GFP-RFP-LC3 were seeded on 2 mg/ml collagen I (Coll) or plastic (P) under complete or Glc starvation media for 1 day. Cells were fixed and stained with Hoechst 33342 and Phalloidin Alexa Fluor 647. Images were collected with a ZEISS LSM98O Airyscan2 confocal microscope and analysed by Fiji/lmageJ software. Data are presented as mean ± SEM, N=3 independent experiments (the bigger dots represent the mean of individual experiments). Non-significant, Kruskal-Wallis, Dunn’s multiple comparisons test.

**Figure S8.**
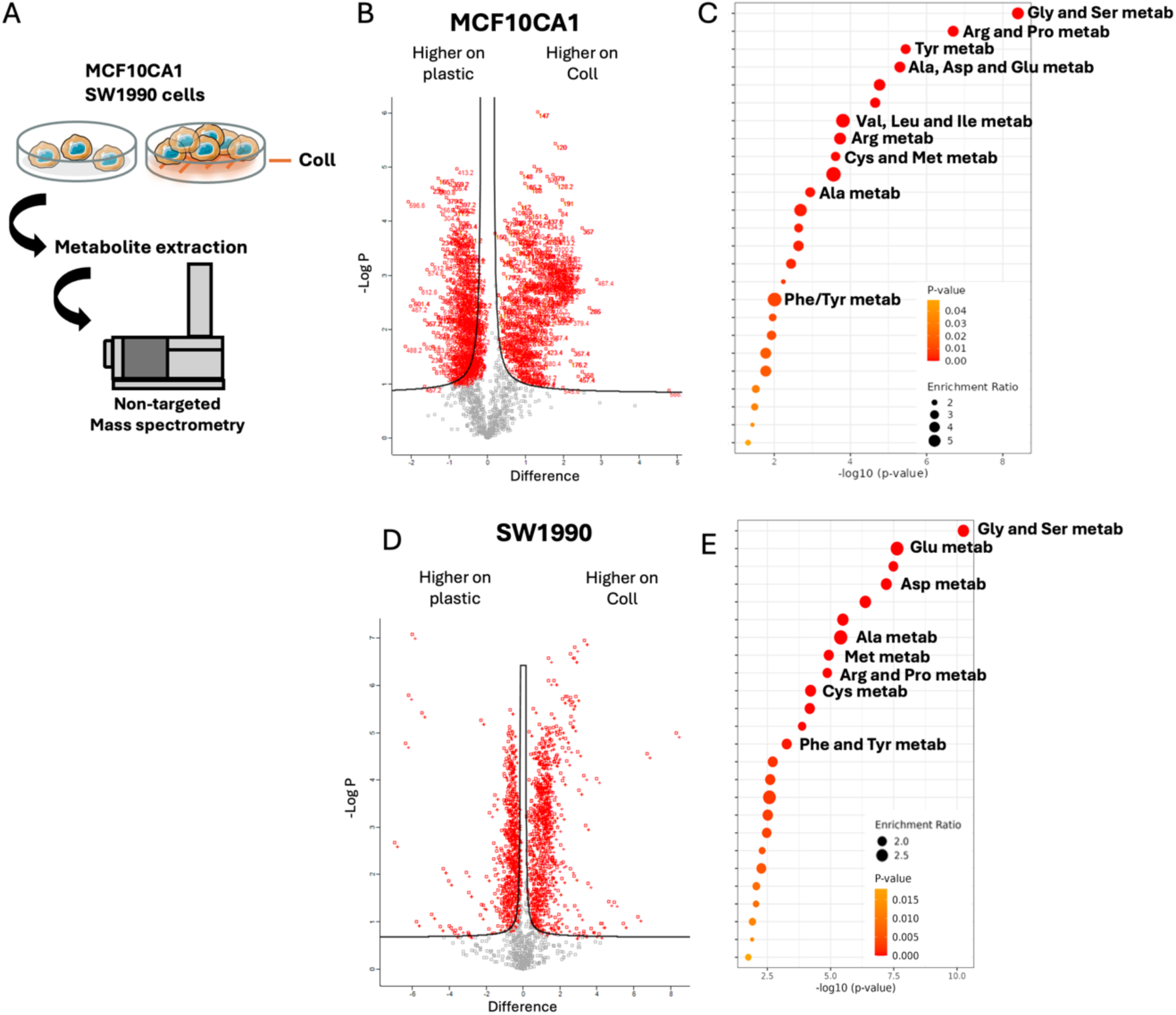
Collagen I increased intracellular amino acid levels in MFC10CA1 and SW1990 cells under glucose starvation. **(A)** Metabolomics workflow. MCF10CA1 and SW1990 cells were plated on plastic or 2 mg/ml collagen I (Coll) in Glc starvation media for 1 day. Metabolites were extracted and quantified by non-targeted mass spectrometry. Volcano plot **(B,D)** and enriched metabolic pathways **(C,E)** are shown.

**Figure S9.**
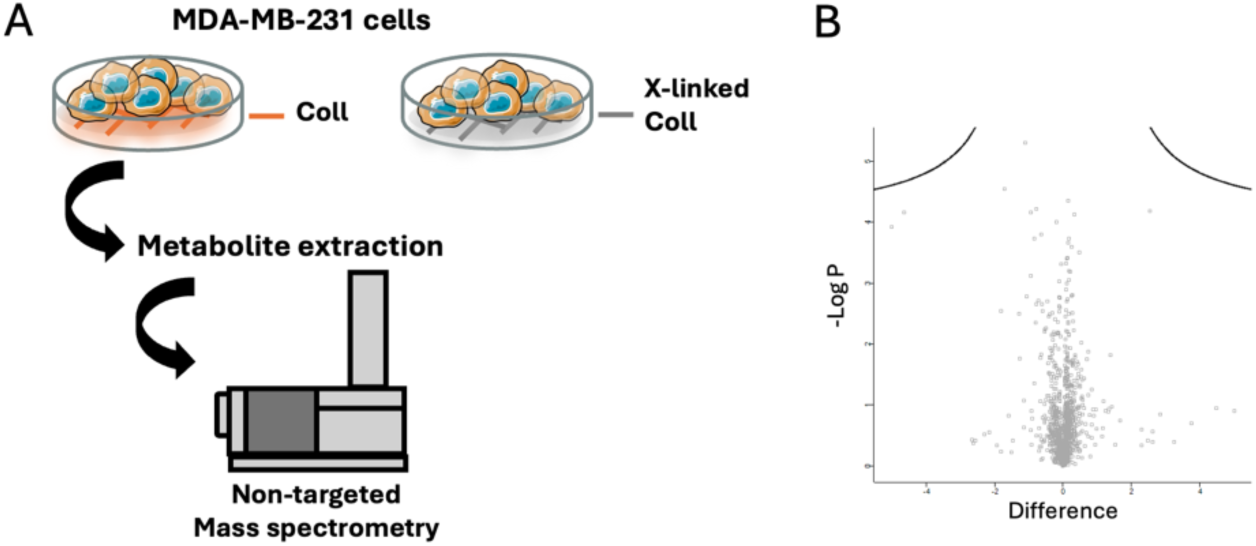
Collagen I crosslinking did not affect intracellular metabolite levels. **(A)** Metabolomics workflow. **(B)** MDA-MB-231 cells were plated on 2 mg/ml collagen I (Coll) or 10% glutaraldehyde cross-linked 2 mg/ml collagen I (X-linked Coll) for 1 day in Glc starvation media. Metabolites were extracted and quantified by non-targeted mass spectrometry. Volcano plot is shown.

